# EpiFlow: multidimensional single-cell epigenetic profiling by spectral flow cytometry

**DOI:** 10.64898/2026.03.16.711606

**Authors:** Jose Ruiz-Iglesias, Elena R. Bovolenta, Lucía Cañizares-Moscato, Javier Isoler-Alcaraz, Lucía Martín-Rodriguez, Joana Segura, Violeta Enriquez-Zarralanga, Adrián de Rus-Moreno, Aida Contreras-Pérez, Alicia Gómez-Moya, Dafne García-Mateos, José Antonio Tercero, Bénédicte Desvoyes, Ana Martínez-Val, Emilio Camafeita, Jesús Vázquez, Daniel Jimenez-Carretero, Berta Raposo Ponce, María Dolores Ledesma, Natalia Reglero, Carlos Perea, Emilio Lecona, María Gómez, Nuria Martínez-Martín, Ernest Palomer

## Abstract

The epigenetic landscape of individual cells determines their identity and function, yet current methods for profiling chromatin modifications at single-cell resolution remain low-throughput, costly, or limited in parametric depth. Here we present EpiFlow, a spectral flow cytometry-based platform that enables the simultaneous quantification of 16 epigenetic markers, including histone post-translational modifications, DNA methylation, and hydroxymethylation, at the single-cell level. We demonstrate that EpiFlow is robust across species from yeast to mammals and resolves biologically meaningful epigenetic transitions during the cell cycle, stem cell differentiation, germinal centre B cell maturation, diabetic liver remodelling, and seizure-induced chromatin reprogramming. High-dimensional integration of EpiFlow data enables cell-type classification based solely on epigenetic profiles in liver, brain, blood, and cancer. Furthermore, EpiFlow detects on-target and off-target/indirect effects of epigenetic drugs in a high-throughput-compatible format. Collectively, these results establish EpiFlow as a broadly applicable platform for single-cell epigenetic analysis in basic, pharmaceutical, and translational research.

## INTRODUCTION

Epigenetic modifications, including DNA methylation and histone post-translational modifications (PTMs), are fundamental in regulating chromatin architecture and gene expression, thereby influencing cellular identity, development, and disease progression^1^. A key challenge in epigenetic research is that these modifications are highly heterogeneous at the single-cell level: even within a seemingly homogeneous population, individual cells can harbour distinct epigenetic states that drive divergent functional outcomes. This heterogeneity is particularly relevant in disease contexts such as cancer, where subpopulations with distinct epigenetic landscapes may have functional consequences^2^. These populations are critical for disease development and treatment, and yet they are invisible to bulk analyses^2^. Traditional methods for analysing epigenetic modifications include molecular biology approaches such as Bisulfite Sequencing (BS-Seq), Methylated DNA immunoprecipitation (MeDIP), chromatin immunoprecipitation sequencing (ChIP), and CUT&Tag, or mass spectrometry for histone PTMs, which have significantly advanced our understanding of epigenetic landscapes^3–5^. However, molecular approaches have limited throughput, involve complex, time-consuming workflows, and allow analysis of only up to two marks simultaneously across the genome^4,5^. In addition, these techniques can be applied at the single-cell level only in particular cases^4^. On the other hand, while mass spectrometry has the power to characterise the full histome PTMs, it generally requires a substantial amount of sample, and has limited throughput^3^. Nonetheless, recent advances in the field have enabled obtaining insights into the single-cell epigenetic profile of histones, although it remains challenging and requires highly specialised facilities^6^. Thus, there is an urgent need to develop new technologies with high content capacity for studies where the sample size is constrained and/or cellular heterogeneity is critical.

The development of EpiTOF, a mass cytometry (CyTOF) based approach, enabled multiplexed histone modification profiling at the single-cell level^7^. However, mass cytometry remains costly due to the need for metal-conjugated antibodies, and its application is severely limited by instrument availability. To overcome these limitations, we have developed EpiFlow, a spectral flow cytometry-based approach that enables the simultaneous measurement of 16 epigenetic marks at the single-cell level. Unlike conventional flow cytometry, which is constrained by spectral overlap between fluorophores, spectral flow cytometry captures the full emission spectrum of each fluorophore and uses computational unmixing to resolve highly overlapping signals, enabling a far greater number of parameters to be measured simultaneously without sacrificing data quality^8^. This multiplexing capacity confers EpiFlow the flexibility and scalability necessary to generate epigenetic profiles in different biological and pathological settings. In addition, spectral cytometry is increasing its presence in research centres, which, coupled with in-house antibody conjugation and the increasing availability of dyes for spectral cytometry, democratises the use of EpiFlow and would allow the study of virtually any epigenetic mark.

In this study, we demonstrate EpiFlow’s robustness and applicability across species, its potential for drug screening, and its capacity to resolve epigenetic dynamics across cell cycle progression, stem cell differentiation, immune cell differentiation, metabolic disease, and status epilepticus. Together, these applications showcase the versatility of EpiFlow and demonstrate its capacity to confirm known epigenetic changes as well as to reveal previously unreported epigenetic modifications at single-cell resolution across diverse experimental contexts. Altogether, EpiFlow offers a flexible and versatile approach to gain an unprecedented view into single-cell epigenetic landscapes, based on an affordable method with broad applicability across multiple biological disciplines.

## RESULTS

### EpiFlow setup and validation

The basic EpiFlow design includes the analysis of total histone levels and 15 epigenetic marks: 12 histone tail modifications (tri-methylation of the lysine 4 of histone H3 (H3K4me3), acetylation of the lysine 9 of histone H3 (H3K9ac), H3K9me3, H3K14ac, H3K27ac, H3K27me3, H3K36me2, H3K36me3, H4K8ac, H4K16ac, H4K20me2, and H4K20me3), one histone core modification (H3K79me3) and 2 DNA modifications (5-mC and 5-hmC; Fig. 1A)^9^. The panel measures: marks enriched in euchromatin and linked to transcriptional activation (H3K9ac, H3K14ac, H3K27ac, H4K8ac, and H4K16ac); marks enriched in heterochromatin and linked to gene repression (H3K9me3, H3K27me3, H4K20me3, 5-mC); marks enriched in promoters and linked to transcriptional initiation (H3K4me3); and marks enriched at gene bodies and linked to transcriptional elongation (H3K36me2, H3K36me3, H3K79me3, 5-hmC; Fig. 1A)^1,9^. We conjugated all the antibodies to spectral-compatible dyes, systematically titrated them (see Methods), and used them first in human samples (*Homo sapiens*; Jurkat; Fig. 1B, gating in Fig. S1A). Note that in the bubble plots, both colour and size indicate abundance. Fluorescent-minus-one (FMO) experiments (Fig. 1B) demonstrate that EpiFlow can be used in human samples with specific signals for each epigenetic mark.

**Figure 1:**
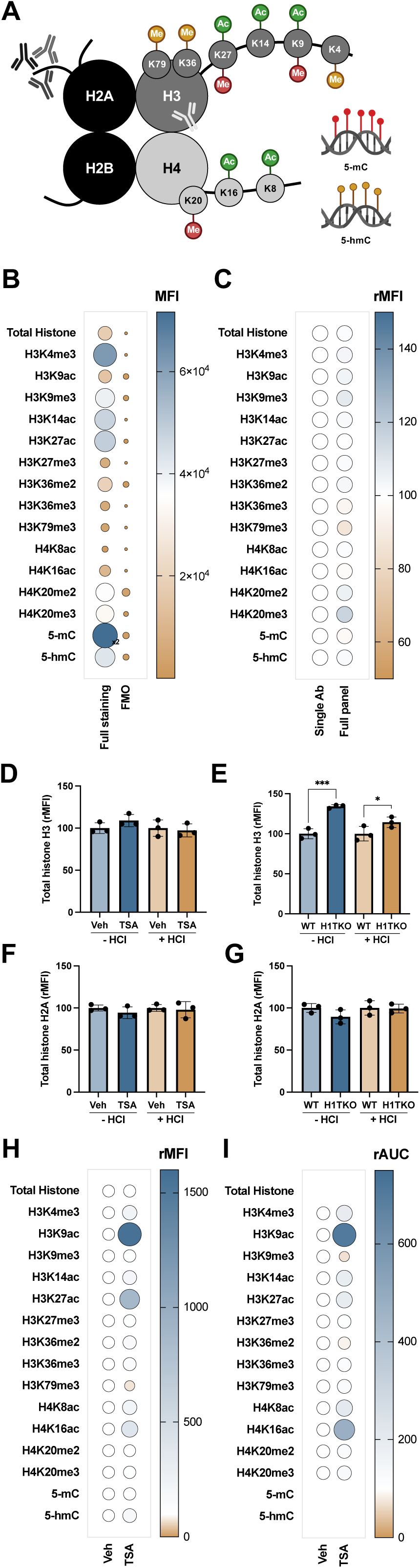
Setting up EpiFlow. **A)** Schematic representation of EpiFlow targets: acetylation (green), heterochromatin-asssociated marks (red) and non-repressive methyl marks and DNA hydroxymethylation (ochre). Antibodies represent monoclonal antibodies against histone H3 core (pale grey) and polyclonal antibodies against the tail of histone H2A (dark grey range). **B)** Bubble plot showing that all antibodies of the EpiFlow panel can be detected (left column, full staining) and that their levels of detection are above their respective FMO (right column, FMO). Note that the FMO column only shows the value for the antibody not present in the mix. Remaining signals for each antibody in each FMO experiment are not shown. The x2 symbol indicates that the 5-mC MFI value has been halved for display purposes and should be multiplied by 2 to obtain the real signal. **C)** Bubble plot showing no allosteric impediments in EpiFlow expressed as the % of relative mean fluorescence intensity (rMFI) for single ab (white; N=3). **D-G)** Bar graphs showing total histone detection in vehicle (Veh)- and TSA-treated Jurkat (D for H3 and F for H2A), and WT and H1TKO mES cells (E for H3 and G for H2A), in the presence or absence of HCl. **H-I)** Bubble plot showing EpiFlow (H N=4) and mass spectrometry (I, N=2-3) results for vehicle (Veh)- and TSA-treated Jurkat expressed as % of Veh (white) by the relative area under the curve (rAUC). Note that both, size and colour represent antibody levels in the bubble plots.

Multiplexing raises two relevant technical questions. First, the use of up to 9 antibodies that bind to rather consecutive amino acids in a single histone could result in allosteric impediments. To rule out allosteric hindrance, we compared the signal of each antibody when used individually versus the signal in the presence of the complete non-fluorescent antibody panel. No significant differences were observed for any of the marks analysed as relative mean fluorescence intensity (rMFI; Fig. 1C). Second, we studied the EpiFlow capacity to detect total histone levels using a monoclonal antibody targeting the histone H3 core domain (immunogen located distally to all PTMs included in the panel; Fig. 1A). This design prevents competition with the other antibodies in the panel targeting PTMs, though the core location may reduce antibody accessibility. To test whether this antibody can access chromatin and reliably measure total histone, we compared untreated chromatin with chromatin relaxed by Trichostatin A (TSA) treatment in Jurkat cells. This epigenetic drug is a pan-histone deacetylase inhibitor that upregulates histone acetylation, thereby relaxing chromatin conformation without altering total histone levels^10^. Total levels of histone H3 remained unchanged upon TSA treatment (Fig. 1D), suggesting that the detection of total histone levels is not conditioned by chromatin condensation. Next, we compared WT mouse embryonic stem (mES) cells and mES cells deficient in histone H1 (H1TKO), which naturally exhibit a relaxed chromatin conformation without altering the levels of core histones^11^. In this case, we detected higher levels of total histone H3 in H1TKO cells compared to WT (Fig. 1E). Given that neither TSA treatment nor H1 deficiency increase the total amount of histone H3^10,11^, yet the detection of histone H3 was elevated in mES cells with more relaxed chromatin, we decided to apply an antigen-retrieval protocol, treating the cells with HCl to denature and even break DNA molecules, thereby allowing better antibody access to their epitopes in fixed cells^12^. HCl treatment reduced the difference in histone H3 detection in mES H1TKO cells, whereas it had no effect on TSA treatment (Fig. 1D-E). As an alternative to the detection of core histone H3, we conducted parallel experiments using a polyclonal antibody against the histone H2A tail (Fig. 1A), which protrudes from the nucleosome core. Further, using a polyclonal antibody reduces the probability of detecting changes in total histone H2A due to the underlying epigenetic changes in this region. Importantly, neither TSA treatment in Jurkat cells nor mES cells with reduced H1 resulted in a significant change in total histone H2A levels, regardless of HCl treatment (Fig. 1F-G). These results support the use of the tail of histone H2A to quantify total histone levels.

Finally, we compared EpiFlow with mass spectrometry detection of changes in epigenetic marks in Jurkat cells treated with TSA (Fig. 1H-I; gating in Fig. S1B). As expected, TSA induced a dramatic increase in histone acetylation (Fig. 1H-I), detected in both by flow cytometry (Fig. 1H) and mass spectrometry (Fig. 1I). These results further confirm the applicability of EpiFlow to properly detect epigenetic changes, validating the use of our basic EpiFlow panel with histone H2A as a marker to quantify total histone levels.

### EpiFlow applicability across species

Histone proteins, their post-translational modifications (Fig. S1C) and DNA modifications are highly conserved across mammals^9^. Therefore, we investigated the applicability of EpiFlow in cells of different mammalian species: monkey (*Chlorocebus aethiops*; COS7), dog (*Canis lupus familiaris*; MDCK), rat (*Rattus norvegicus*; PC12), mouse (*Mus musculus*; N2A), and human (*Homo sapiens*, Jurkat, Fig. 2A, cell line gating in Fig. S1A). Our results show that EpiFlow can indeed be extended to study epigenetics across mammalian species (Fig. 2A). In addition, histones and their PTMs are conserved across eukaryotic organisms (Fig. S1C), whereas this is not the case for DNA modifications, which are absent or present at low levels in non-mammalian cells^9,13–15^. Thus, we tested EpiFlow in insects (*Trichoplusia ni*; Hi5; Fig. 2B, cell line gating in Fig. S1A), plants (C2 root of *Arabidopsis thaliana*, Fig. 2C, gating in Fig. S1D) and yeast (*Saccharomyces cerevisiae*, Fig. 2E, gating in Fig. S1E). We demonstrated that EpiFlow can be successfully applied across the whole range of eukaryotic species, highlighting its robustness and broad potential for comparative epigenetic studies and evolutionary biology research.

**Figure 2:**
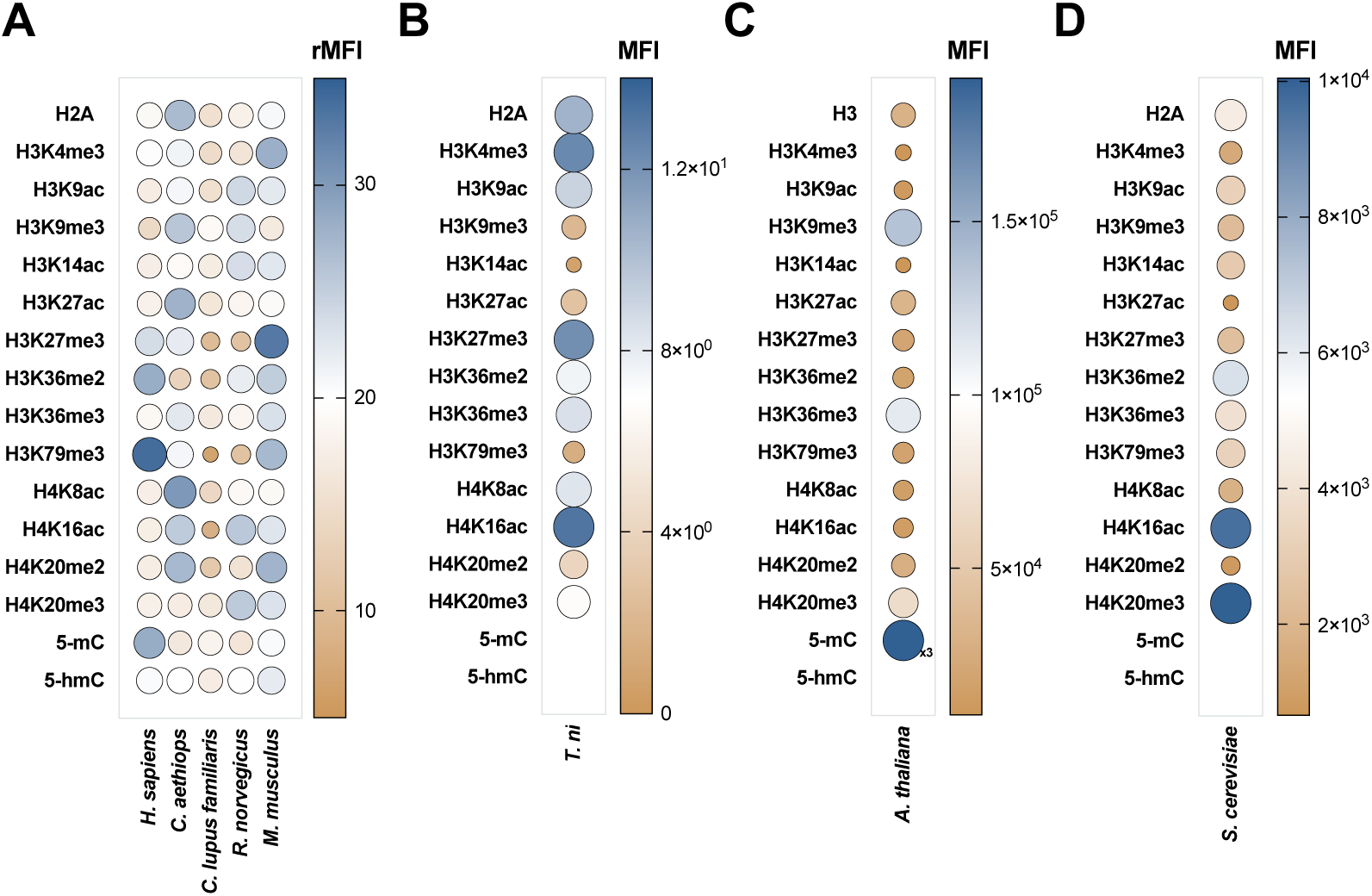
EpiFlow across eukaryotes. **A-D)** Bubbe plots showing EpiFlow across eukaryotes: mammals (A; *Homo sapiens*, Jurkat; *Chlorocebus aethiops*, COS7; *Canis lupus familiaris*, MDCK; *Rattus norvegicus,* PC12; *Mus musculus*; N2A), insects (B; *Trichoplusia n*i, Hi5), plants (C; C2 root of *Arabidopsis thaliana*) and yeast (D; *Saccharomyces cerevisiae*). Data are represented as the mean of biological replicates (N=4). In C, the x3 symbol indicates that the 5-mC MFI value is displayed at one-third scale for display purposes and should be multiplied by 3 to obtain the real signal. Gating strategies, shown in Fig. S1.

### Analysis of drug-induced epigenetic changes using EpiFlow

Next, we wanted to evaluate whether EpiFlow can detect alterations induced by drugs targeting epigenetic modulators. The different epigenetic marks included in the EpiFlow panel are enriched in specific regions of the genome, either linked to transcription activation or repression, or specifically involved in transcriptional initiation or elongation, or with a role in chromatin structure. To facilitate the rapid visualisation of epigenetic landscapes and their changes upon treatment, we decided to integrate single-cell data into scores with biological meaning: Chromatin relaxation (∑ Histone acetylation/∑ Heterochromatin marks; CR), DNA Methylation (5-hmC/5-mC; DNA Meth), Facultative Heterochromatin Activation (H3K27ac/H3K27me3; FHcA), Facultative vs Constitutive Heterochromatin (H3K27me3/H3K9me3; FHc/CHc), Heterochromatin Index (∑ Heterochromatin marks; HcI), Enhancer activity (H3K27ac/H3K4me3; EA), Transcription Initiation and Elongation (H3K4me3 + H3K36me2 + H3K36me3 + H3K79me3 + 5hmC; I&E), Transcriptional Potential (∑ activation marks/∑ repressive marks; TP, Fig. S1F, see Methods). As a proof of concept, we first treated Jurkat cells with TSA and followed the changes in epigenetic marks normalised against total histone levels (marked with an “n” in front of the histone mark, see Methods). As expected, our results showed an increase in histone acetylation upon TSA treatment, along with changes in other marks such as an increase in H3K4me3 or a decrease in H3K79m3 (Fig. 3A, gating in Fig. S1B). The multidimensional epigenetic space integration by EpiFlow scores reflected an increase in chromatin relaxation and transcriptional activation due to increased histone acetylation, with no change in scores composed solely of heterochromatin marks (Fig. 3B). Thus, EpiFlow allows not only the detection of on-target effects of epigenetic drugs, but also their off-target/secondary effects in a single experiment.

**Figure 3:**
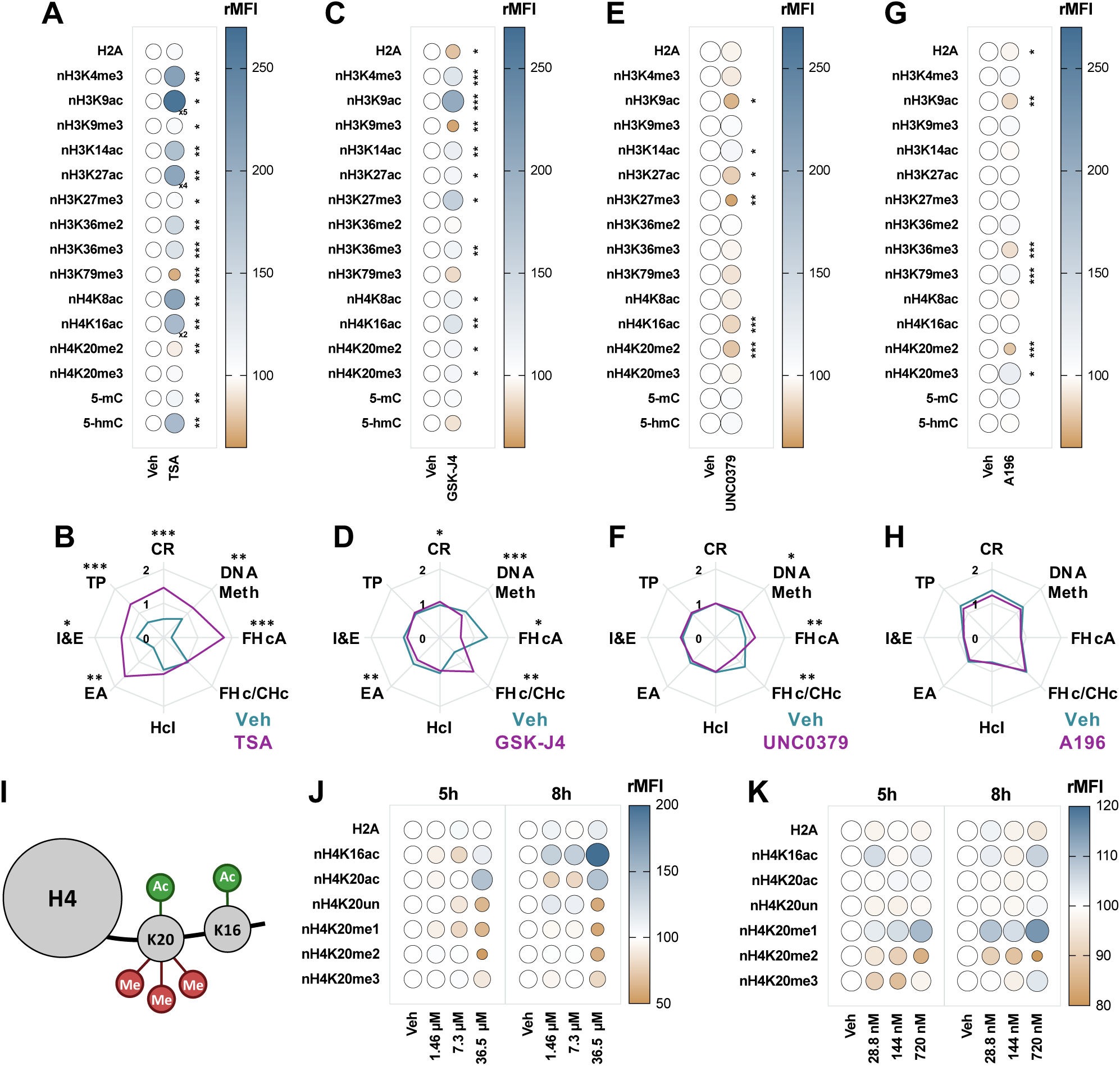
EpiFlow to study drug-induced epigenetic changes. **A-H)** Bubble and spider plots showing epigenetic changes induced by TSA (5h; A-B), GSK-J4 (24h; C-D), UNC0379 (5h; E-F) and A196 (5h; G-H). In A, the x2, x4 and x5 symbols indicate that the respective MFI values have been rescaled for display purposes and should be multiplied accordingly to obtain the true signal. **I)** Schematic representation of the alternative EpiFlow panel used for a high-content approach. **J-K)** Bubble plots showing epigenetic changes induced by increasing concentrations of UNC0379 (J) and A196 (K) at two time points, 5h and 8h (N=4). All experiments were performed in Jurkat cells. Data are shown as the mean of biological replicates and expressed as % of Veh (white; N=4). Gating strategies, shown in Fig. S1. Statistical analysis was performed using Student’s t-test (A-H). *p□<□0.05; **p□<□0.01, ***p□<□0.001. Statistics for I-K are provided in Table S1. Note that histone marks precede by “n” indicate that the levels represented are normalised by the total histone levels.

To further test the capacity of EpiFlow to follow epigenetic changes, we treated Jurkat cells with three more specific inhibitors: we targeted the H3K27me3 demethylases KDM6A/B with GSK-J4^16^ (Fig. 3C-D); H4K20me1 methyltransferase KMT5A with UNC0379^17^ (Fig. 3E-F); and H4K20me2/3 methyltransferases KMT5B/C with A196^18^ (Fig. 3G-H). In all cases, we detected the expected changes in the epigenetic marks targeted by these enzymes. Additional off-target/secondary effects were observed, leading to changes in specific EpiFlow scores. Upon treatment with GSK-J4, we observed increased H3K27me3 levels, as well as increased H3K4me3 (Fig. 3C), a known off-target effect of the drug^16^. Accordingly, GSK-J4 induced changes in facultative heterochromatin scores, related to H3K27me3 (Fig. 3D). Both UNC0379 (Fig. 3E-F) and A196 (Fig. 3G-H) led to reduced levels of H4K20 methylation. While A196 did not change the epigenetic landscape (Figure 3H), UNC0379 also led to changes in facultative heterochromatin (Figure 3F). Altogether, these results prove that EpiFlow detects the effects of epigenetic drugs and that the EpiFlow scores allow capturing the drug-induced changes in epigenetic landscapes.

Finally, to evaluate the application of EpiFlow for high-content screening (HCS) of epigenetic drugs, we carried out dose and time-dependent analysis of the effects of UNC0379 or A196 in Jurkat cells. For this experiment, we developed an H4K20-focused EpiFlow panel, including acetylation (H4K20ac), unmodified (H4K20un), the three methylation states (H4K20me1/2/3), and H4K16ac, known to be inversely co-regulated with H4K20me (Fig. 3I)^19^. Targeting the H4K20me1 methyltransferase KMT5A with UNC0379 effectively reduced H4K20me1 levels in a dose- and time-dependent manner (Fig. 3J), leading to a progressive decrease in H4K20me2/3 levels and a concomitant increase in H4K20ac and H4K16ac (Fig. 3J). A196 treatment induced the reduction of H4K20me2 and the consequent accumulation of H4K20me1, the substrate of KMT5B/C (Fig. 3K). No large effect on other marks at the dosages and treatment times used (Fig. 3K). These results demonstrate EpiFlow’s scalability for high-throughput applications and its applicability for drug screening, detecting both direct effects and indirect consequences (off-target/secondary effects) of epigenetic modulators.

### EpiFlow in biological settings used in research

After determining the capacity of EpiFlow to detect drug-induced epigenetic changes in cultured cells, we tested its capacity to identify epigenetic changes across different physiological and pathological settings, from cell-cycle and mES de-differentiation to disease.

#### Cell cycle

To assess whether EpiFlow can resolve the cell-cycle dynamics of epigenetic marks, we analysed asynchronous HeLa-S3 cells. Cell cycle phases were determined using EdU incorporation combined with 7-AAD DNA content, together resolving G1, early S (eS), middle (mS) and late (lS), and G2 phases (for gating see Fig. S2A). Our results show that total histone levels, measured by H2A, increase through S phase and G2 (Fig. 4A), consistent with coupled histone deposition at the replication fork^20^. Active histone marks, including H3K4me3, H3K9ac, H3K14ac, H3K27ac, H3K36me3, H3K79me3, increased through S phase (Fig. 4A), consistent with their fast restoration on newly synthesised chromatin^20^. As expected, 5-mC levels follow a similar pattern, consistent with DNMT1-mediated maintenance of DNA methylation, directly linked to the replication machinery at the fork (Fig. 4A)^21^. 5-hmC remained unchanged throughout S and G2 phases (Fig. 4A), as it is generated post-replicatively by TET-mediated oxidation of newly DNMT1-methylated cytosines^22^. A similar result was obtained for H4K8ac and H4K16ac (Fig. 4A), reflecting the incorporation of newly deposited H4 carrying the CAF-1 deposition signature (H4K5ac/K12ac) rather than H4K8ac and H4K16ac, which are acquired post-replication^23^. Repressive heterochromatin marks were restored more slowly as cells progress through the cell cycle (Fig. 4A). For example, H3K27me3 was increased during late S phase and G2, consistent with PRC2 uncoupling from the replication machinery and its dependence on allosteric re-stimulation by pre-existing H3K27me3 on parental histones for inheritance^24^. Similarly, the levels of H4K20me2 and H4K20me3 increased in late S phase, reflecting the progressive nature of KMT5B/C-mediated re-methylation on newly deposited H4K20 (Fig. 4A)^20^. Together, these results show that EpiFlow allows precise capturing of epigenetic marks deposition throughout the cell cycle. Importantly, using EpiFlow to study epigenetic marks dispositioning in asynchronous cultures will prevent possible artefacts coming from synchronisation protocols.

**Figure 4:**
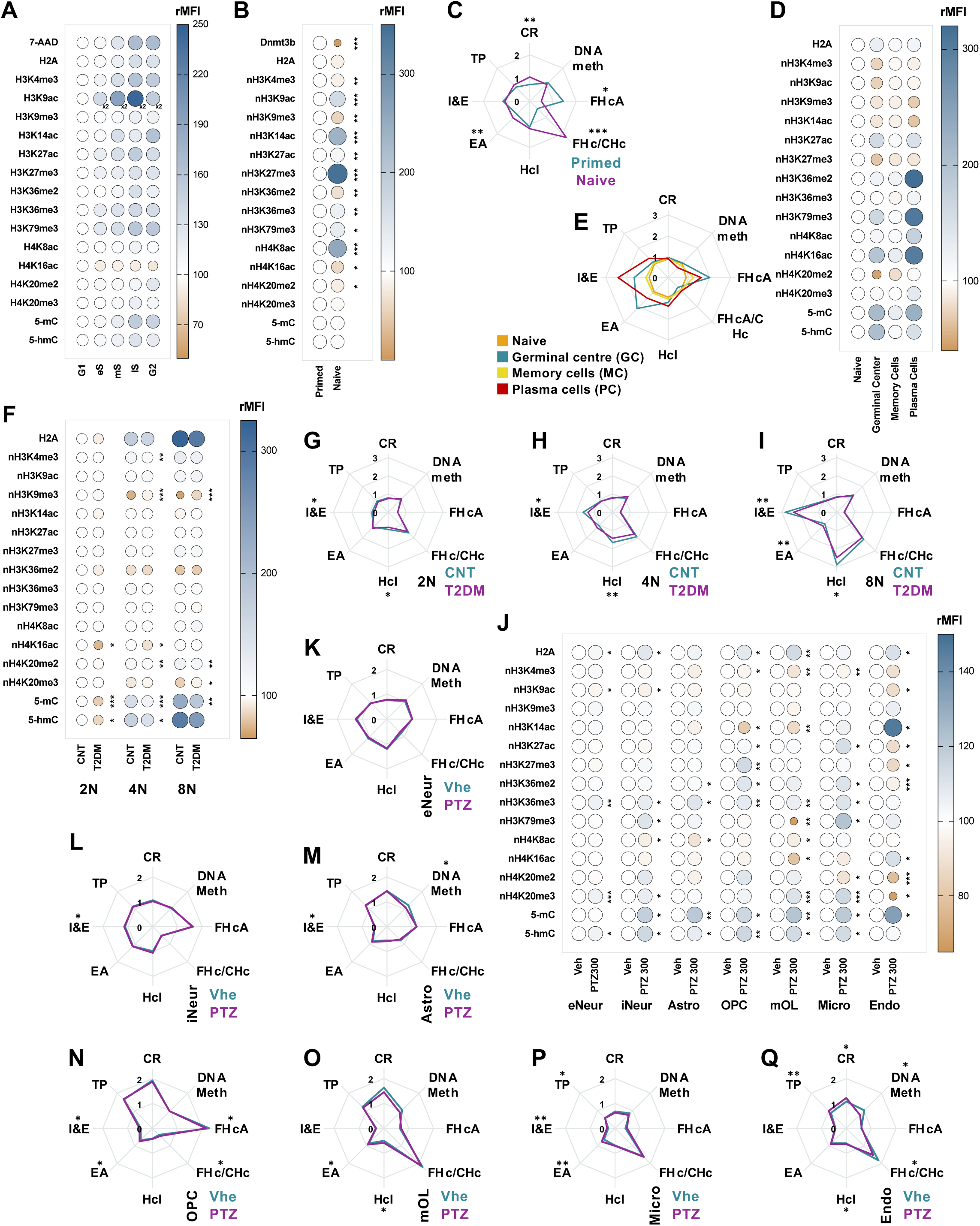
EpiFlow in biological settings used in research. **A)** Bubble plot showing the variation of different epigenetic marks across the cell cycle: G1, early S (eS), middle S (mS), late S (lS), and G2. Data expressed as % of G1 (white; N=4). The x2 symbol indicates that the H3K9ac MFI values have been halved for display purposes and should be multiplied by 2 to obtain the true signal. **B-C)** Bubble plot (B) and spider plot (C) showing epigenetic remodelling during the transition of primed (cyan) to naïve (magenta) mouse embryonic stem cells. Data expressed as % of primed (white; N=3). **D-E)** Bubble plot (D) and spider plot (E) showing the epigenetic remodelling of splenic B cells differentiation during humoral immune response. Data expressed as % of Naive (white; N=7). Naive (orange), germinal centre (GC, cyan), memory cells (MC, yellow) and plasma cells (PC, red). **F-I)** Bubble plot (F) and spider plots (G-I) showing the epigenetic remodelling induced by T2DM in hepatocytes. Control (CNT, cyan) and T2DM (magenta) for hepatocytes 2N (G), 4N (H) and 8N (I). Data expressed as % of CNT 2N (white) in F. N=5-6. **J-Q)** Bubble plot (J) and spider plots (K-Q) showing the seizure-induced epigenetic remodelling in status epilepticus (N=5). Data expressed as % of Veh (white) in J. Control (CNT, cyan) and status epilepticus (PTZ, magenta) for excitatory neurons (eNeur, K), inhibitory neurons (iNeur, L), astrocytes (Astro, M), oligodendrocyte precursor cells (OPC, N), mature oligodendrocytes (mOL, O), microglia (Micro, P) and endothelial cells (Endo, Q). Data are shown as the mean of biological replicates. Gating strategies are shown in Fig. S2. Statistical analysis was performed using Student’s t-test (B-C and F-Q). *p□<□0.05; **p□<□0.01, ***p□<□0.001. Statistics for A and D-E are provided in Table S2. Note that histone marks precede by “n” indicate that the levels represented are normalised by the total histone levels.

#### mES de-differentiation

We characterised the epigenetic remodelling that accompanies the transition from primed to naïve pluripotency using EpiFlow. mES cells were cultured in 2i medium for 7 days. Cells at Day 0 and Day 7 were then barcoded, mixed, and subjected to full EpiFlow staining, enabling direct comparison of two de-differentiation time points in a single tube (see Methods, Fig. S2B). DNA methyltransferase 3b (Dnmt3b) was included as a positive control, given its well-established downregulation during the primed-to-naïve transition^25^. Our results confirmed previously reported epigenetic changes, including a marked increase in H3K27me3 and a reduction in H3K9me3^26^. Further, we observed a general increase in histone acetylation (Fig. 4B), an effect not previously reported. these modifications were integrated into multidimensional epigenetic space, the most prominent shifts between primed and naïve cells were observed in heterochromatin-associated scores (Fig. 4C). This shift in heterochromatin balance is consistent with the notion that primed pluripotent cells exhibit a more lineage-committed chromatin architecture, with greater constitutive heterochromatin consolidation through H3K9me3, whereas naïve cells rely more heavily on H3K27me3-based facultative heterochromatin to silence these same domains, reflecting a more plastic and developmentally open chromatin configuration^26,27^.

#### B cell differentiation in humoral immune response

To profile the epigenetic dynamics along the humoral immune response, mice were immunised with sheep red blood cells (SRBC) and splenic B cells were analysed at day 7 post-challenge. We profiled naive B cells, germinal centre cells (GCt), memory cells (MC), and plasma cells (PC; Fig. 4D-E; surface markers for cell identification and gating strategy see Fig. S2C) using EpiFlow. GCt cells showed a decrease in H3K27me3 with a reciprocal gain in H3K27ac, consistent with redistribution of H3K27me3 to specific bivalent loci^28^, and increased H3K27 acetylation (Fig. 4D) that activates GC-specific enhancers and superenhancers^29^. GCt cells also displayed a simultaneous increase in 5-mC and 5-hmC, consistent with the increased methylome diversity characteristic of GC B cells^30^. Memory cells largely reverted to a naive-like epigenetic profile (Fig. 4D), consistent with the model that MCs reset GC-associated chromatin modifications while retaining subtle epigenetic priming for rapid reactivation^31,32^. MCs retained modestly elevated 5-mC levels (Fig. 4D), which could be part of the progressive DNA methylation changes acquired during GC transit and inherited through memory differentiation^30^. PC exhibited the most dramatic remodelling, dominated by large increases in active transcription and initiation/elongation-associated marks (Fig. 4D). PCs also displayed pronounced 5-mC hypermethylation, consistent with *de novo* methylation programs that consolidate silencing of the B cell identity program during terminal differentiation^33,34^. The decrease in H3K27me3 in PCs is consistent with the removal of Polycomb-mediated repression at plasma cell determinants^33^. The spider plot visualisation of the integrated EpiFlow scores showed that each differentiation stage occupies a distinct multidimensional epigenetic space (Fig. 4E). GCt are defined by reciprocal H3K27me3/H3K27ac remodelling, MCs by the reversion to a naive-like state, and PCs by their increase in transcription activation-related scores. Together, these data demonstrate that EpiFlow differentiates the global epigenetic identity at each differentiation stage of the humoral response and at single-cell resolution, capturing the epigenetic landscape associated to the chromatin dynamics at these stages.

#### Type 2 Diabetes Mellitus in the liver

To define the epigenetic remodelling in the liver in Type 2 Diabetes Mellitus (T2DM), we applied EpiFlow to profile nuclear samples from hepatocytes stratified by ploidy status (2N, 4N, and 8N, gating in Fig. S2D) from control (CNT) and T2DM mice. Our results showed ploidy-dependent epigenetic differences between control and diabetic hepatocytes, with notable changes in H3K9me3, H4K16ac, H4K20me3 and DNA methylation (Fig. 4F). The observed retention of higher levels of H4K20me3 in polyploid T2DM hepatocytes compared to control conditions is consistent with recent findings in liver biopsies from human and mice models^35^. Similarly, lower DNA methylation in T2DM hepatocytes also recapitulates previous reports^36^. Multidimensional integration of epigenetic states revealed progressively increased EpiFlow scores for heterochromatin index and transcriptional initiation/elongation in polyploid hepatocytes (Fig. 4G-I). Interestingly, this is due to the cooperative effect of small, not significant, changes in individual histone methylations (Fig. 4F-I). In T2DM, the increase in the heterochromatin index was significantly attenuated (Fig. 4G-I), as expected.

#### Epilepsy

To characterise the effect of epileptic seizures on brain cells epigenetics, we applied EpiFlow in nuclear samples from vehicle- and GABA antagonist pentylenetetrazol (PTZ)-treated mice sacrificed 5 hours post-injection (Fig. 4J-Q). We first confirmed PTZ-induced epilepsy (Fig S2E) and subsequently profiled seven brain cell populations. We analysed excitatory neurons (eNeur), inhibitory neurons (iNeur), astrocytes (Astro), oligodendrocyte precursor cells (OPC), mature oligodendrocytes (mOL), microglia (Micro) and endothelial cells (Endo; nuclear markers for cell identification and gating strategy see Fig. S2F). Overall, glial and vascular populations were the most affected, whereas iNeur showed greater changes than eNeur. Cell-type-resolved data revealed cell-specific changes in different residues, which would be masked in bulk preparations and may reflect the distinct epigenetic signalling in each cell type. We observed decreased H3K9ac and H4K8ac across almost all cell types, while H3K14ac and H4K16ac increased in microglia and endothelial cells, respectively (Fig. 4J). In contrast, H3K27ac increased in microglia and decreased in endothelial cells (Fig. 4J). The most consistent finding across all cell types was an increase in 5-hmC, in line with the previously reported seizure-induced activation of TET-mediated oxidation of 5-mC^37^ (Fig. 4J). The parallel increase in 5-mC (Fig. 4J) also recapitulates previous studies showing increased global DNA methylation in chronic epilepsy^38^, suggesting that the methylation gain observed in bulk studies may not be from excitatory neurons. The role of histone methylation in seizure-induced epigenetic remodelling has remained essentially unstudied. The EpiFlow analysis shows that the epigenetic profiles of glial and vascular cells are more affected in epilepsy than those of neurons (Fig. 4J). As in the case of the humoral immune response, multidimensional epigenetic integration of EpiFlow scores revealed differential cell-type-specific epigenetic landscapes (Fig. 4K-Q). In addition, spider plots showed subtle differences in scores, mostly in glial and vascular cells rather than in neurons (Fig. 4K-Q). These findings demonstrate that EpiFlow can reveal coordinated, cell-type-specific chromatin state transitions in the brain, suggesting that glial and vascular cells, rather than neurons alone, are the main targets of status epilepticus-induced epigenetic reprogramming.

### EpiFlow potential to identify cell heterogeneity in complex samples

Our results showed that the integrated analysis of EpiFlow scores reveals cell-type-specific multidimensional epigenetic spaces. Thus, we tested whether EpiFlow could dissect cell heterogeneity in complex samples using higher-dimensional reductions and unsupervised clustering. To this end, we processed flow data using staining-specific cofactors for Asinh transformation, followed by robust z-score normalisation to centre EpiFlow stainings (see Methods).

#### EpiFlow in naturally occurring complex samples

First, we tested EpiFlow in mouse nuclear liver samples composed of endothelial cells, Kupffer cells, and hepatocytes (2N, 4N and 8N; nuclear markers for cell identification and gating strategy see Fig. S3A). Manual gating of these populations showed that hepatocytes had a different epigenetic profile than Kupffer and endothelial cells, which presented a rather similar pattern (Fig. 5A). Ploidy-specific patterns were very clear in hepatocytes (Fig. 5B), confirming our previous results by multidimensional epigenetic space integration of EpiFlow scores (Fig. 4G-I). This analysis also showed similar space occupancies for Kupffer and endothelial cells (Fig. 5B). Liver cells were clustered in UMAP space using only the epigenetic marks to perform high-dimensional reduction (Fig. 5C). Unsupervised clustering identified five clusters (Fig. 5D), separating 2N hepatocytes from 4N and 8N hepatocytes and grouping Kupffer and endothelial cells into the same cluster as identified by manual gating of the known populations (Fig. 5E). The two additional clusters formed by unidentified cells arose (Fig. 5D-E), which could represent hepatic stellate cells, cholangiocytes, or immune cells in the liver parenchyma.

**Figure 5:**
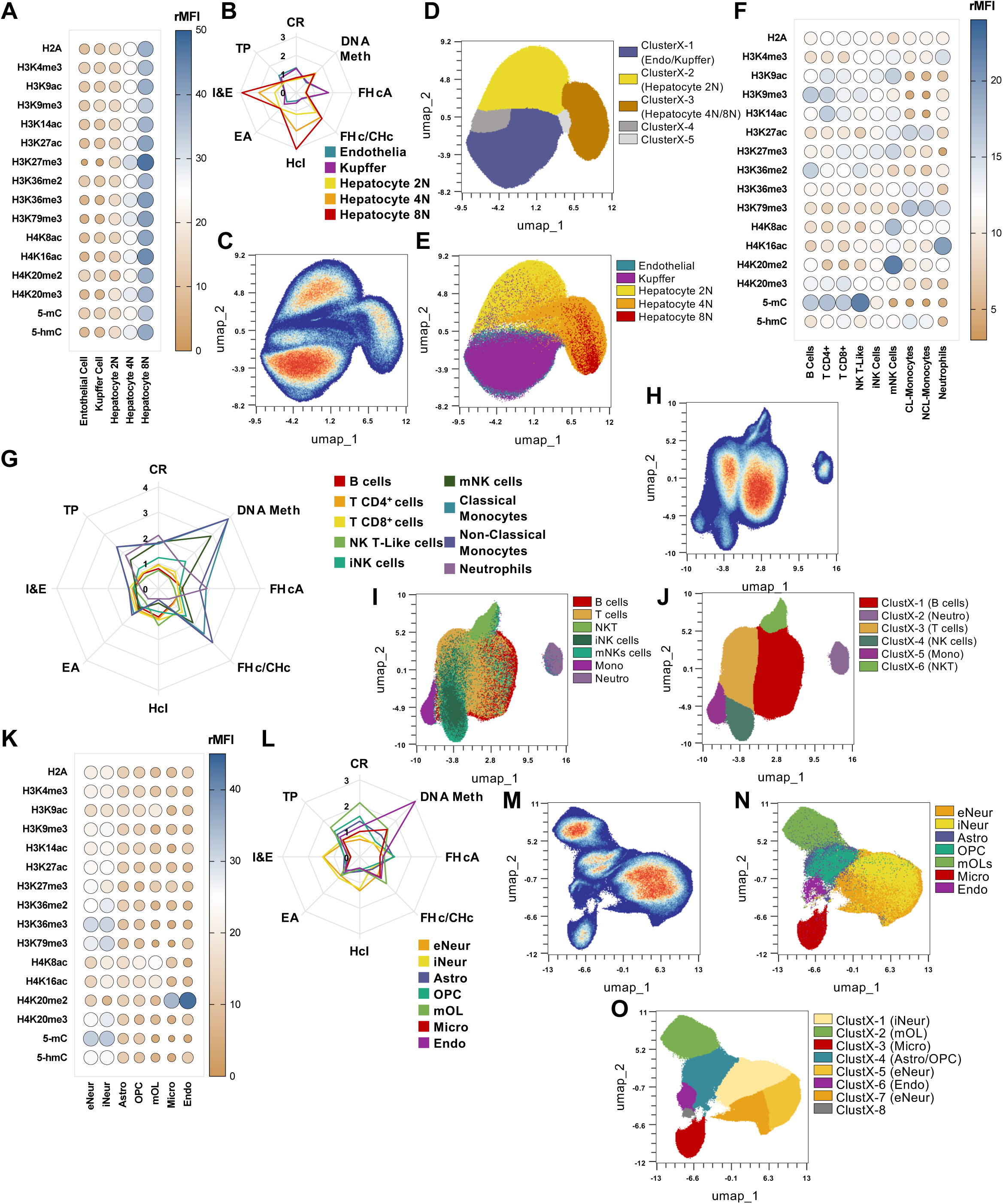
EpiFlow potential to identify cell heterogeneity in naturally occurring complex samples. **A-E)** Bubble plot (A), spider plot (B) and UMAPs projections (C-E) showing cell heterogeneity in mouse liver. Endothelial cells (cyan), Kupffer cells (magenta), hepatocytes 2N (yellow), 4N (orange) and 8N (red). UMAPs display single-cell density (C), manual annotation (D) and unsupervised clustering (E). **F-J)** Bubble plot (F), spider plot (G) and UMAPs (H-J) showing cell heterogeneity in mouse blood. B cells (red), CD4 T cells (orange), CD8 T cells (yellow), natural killer (NK) T-like cells (pale green), immature NK cells (iNK, turquoise green), mature NK cells (mNK, dark green), classical monocytes (CL-Mono, blue), non-classical monocytes (NCL-Mono, dark purple), and neutrophils (Neutro, pale purple). UMAPs show single-cell density (H), manual annotation (I) and unsupervised clustering (J). **K-O)** Bubble plot (K), spider plot (L) and UMAPs (M-O) showing cell heterogeneity in the mouse brain. Excitatory neurons (eNeur, orange), inhibitory neurons (iNeur, yellow), astrocytes (Astro, purple), oligodendrocyte precursor cells (OPC, turquoise green), mature oligodendrocytes (mOL, pale green), microglia (Micro, red) and endothelial cells (Endo, magenta). UMAPs show single-cell density (M), manual annotation (N) and unsupervised clustering (O). Gating strategies shown in Fig. S2-3. Data are shown as the mean of biological replicates (N=4). Statistics are provided in Table S3.

Next, we analysed 9 cell types in murine blood samples: B cells, T CD4 and CD8 cells, Natural Killer T-like (NKT-like) cells, immature NK (iNK) cells, mature NK (mNK) cells, classic and non-classic monocytes (CL-Monocytes and NCL-Monocytes), and neutrophils (surface markers for cell identification and gating strategy see Fig. S3B). Manual gating of these populations showed cell-type-specific patterns (Fig. 5F). When integrated into multidimensional epigenetic spaces by EpiFlow scores, adaptive immune cells occupied similar epigenetic spaces, while innate cells presented specific epigenetic landscapes (Fig. 5G). Interestingly, when only epigenetic marks were used to perform high-dimensional reduction, blood cells clustered in UMAP space (Fig. 5H), yielding 6 individual clusters where we could identify B cells, T cells, NKT-like cells, NK cells, monocytes, and neutrophils (Fig. 5I-J).

Finally, we analysed the 7 mouse brain cell types as per previous experiments (gating in Fig. S2E). Manual gating revealed similar epigenetic profiles in excitatory neurons (eNeur) and inhibitory neurons (iNeur), while glial and endothelial cells showed rather cell-type-specific profiles (Fig. 5K). Multidimensional epigenetic space integration showed similar results, where neurons presented similar epigenetic landscapes while glial and endothelial cells presented cell-type-specific ones (Fig. 5L). Brain samples also clustered in UMAP space when only epigenetic marks were used for high-dimensional reduction (Fig. 5M). Comparing manual annotation of known cell-type populations to unsupervised clustering revealed that EpiFlow data allowed discrimination of microglia, oligodendrocytes, and endothelial cells as single clusters (Fig. 5N-O). We also identified three neuronal clusters, one being inhibitory neurons, but could not differentiate astrocytes from OPCs, even if two different clusters seemed to arise in UMAP space, one belonging to each cell type (Fig. 5N-O). In addition, an extra cluster appeared in the unsupervised clustering, possibly corresponding to other cell types, not analysed in our experiments. We conclude that EpiFlow can dissect cell-type heterogeneity in complex tissues, supporting the existence of cell-type-specific epigenetic profiles that define each cell type.

#### EpiFlow in complex artificial mixtures

Since EpiFlow can distinguish cell heterogeneity in complex tissues, we wondered whether it could identify cell types in artificial mixtures. Endothelial cells exhibit tissue-specific properties^39^, and resident macrophages have tissue-specific functions in the brain (microglia), the lung (alveolar macrophages), and the liver (Kupffer cells)^40^. We explored whether these specific features can be separated using unsupervised algorithms to analyse EpiFlow data. Brain, lung and liver nuclear tissue preparations were barcoded for tissue identification and stained with cell-type markers. Subsequently, samples were mixed, enabling the simultaneous study of endothelial cells and resident macrophages in the same sample by EpiFlow (Fig. 6A, gating in Fig. S4A; see Methods). Manual deconvolution of barcoding and annotation of macrophages revealed tissue-specific epigenetic patterns (Fig. 6B) and distinct integrated multidimensional epigenetic spaces (Fig. 6C) for microglia, alveolar macrophages, and Kupffer cells. In addition, tissue-specific macrophages clustered independently in UMAP space (Fig. 6D), where unsupervised clustering identified two clusters for microglia and specific clusters for alveolar macrophages and Kupffer cells (Fig. 6E-F). These results support the notion that macrophages acquire tissue-specific epigenetic profiles to fulfil their functions. A similar scenario emerged for endothelial cells, which displayed tissue-specific epigenetic patterns (Fig. 6G) and distinct multidimensional epigenetic space occupancies (Fig. 6H). Tissue-specific endothelial cells clustered in two blocks in UMAP space, where liver endothelial cells clustered separately from brain and lung (Fig. 6I). Furthermore, unsupervised clustering identified three clusters for lung endothelium and single clusters for brain and liver endothelial cells (Fig. 6J-K). These results correlate with a tighter barrier in the brain and lung to prevent the entry of exogenous organisms into the brain and the body, and a highly permeable endothelium in the liver to support blood filtration and metabolic processing.

**Figure 6:**
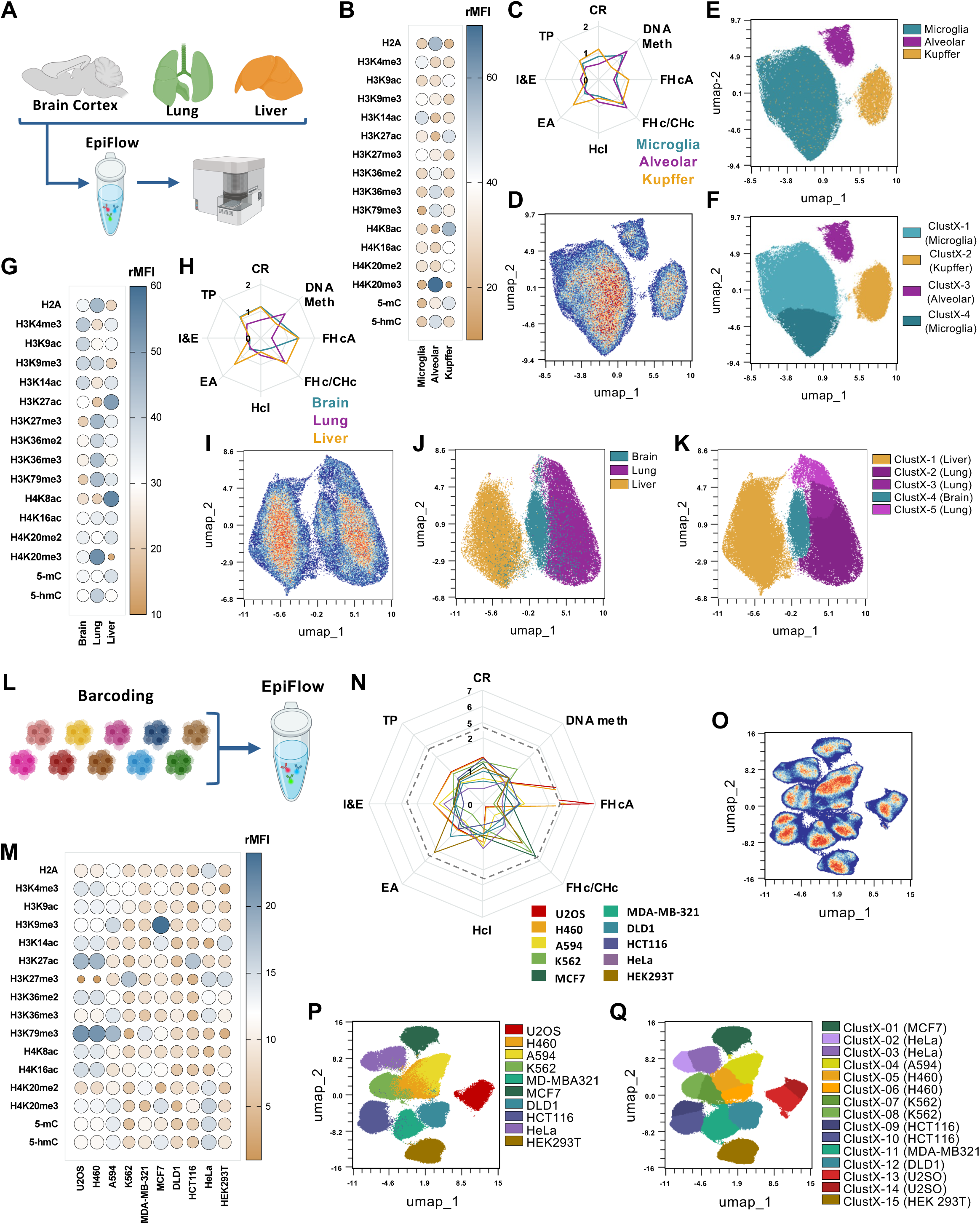
EpiFlow potential to identify cell heterogeneity in artificial mixtures. **A)** Schematic representation of the barcoding staining used for labelling endothelial cells and tissue-resident macrophages in the brain, lung and liver. **B-F)** Bubble plot (B), spider plot (C) and UMAPs (D-F) showing tissue resident macrophage heterogeneity in mouse brain (microglia, cyan), lung (alveolar macrophages, magenta) and liver (Kupffer cells, ochre, N=4). UMAPs display single-cell density (D), manual annotation (E) and unsupervised clustering (F). **G-K)** Bubble plot (G), spider plot (H) and UMAPs (I-K) showing endothelial cell heterogeneity in mouse brain (cyan), lung (magenta) and liver (ochre, N=4). UMAPs display single-cell density (I), manual annotation (J) and unsupervised clustering (K). **L)** Schematic representation of the barcoding staining used to label cancer cell lines. **M-Q)** Bubble plot (M), spider plot (N) and UMAPs (O-Q) showing cell heterogeneity in human-derived cancer cell lines and immortalised HEK293T. Osteosarcoma (U2OS, red), non-small cell lung cancer (H460, orange; A594 yellow), leukaemia (K562, pale green), breast cancer (MCF7, dark green; MDA-MB-231, turquoise green), colorectal carcinoma (DLD1, blue; HCT116, dark purple), cervical cancer (HeLa, pale purple), and HEK293T (brown, N=6). UMAPs display single-cell density (O), manual annotation (P) and unsupervised clustering (Q). Gating strategies shown in Fig. S4. Data are shown as the mean of biological replicates. Statistics are provided in Table S4.

Finally, we tested whether EpiFlow can tell apart epigenetic heterogeneity in cancer cell lines. We analysed 10 cell lines of diverse origins, including osteosarcoma (U2OS), non-small cell lung cancer (H460, A594), chronic myelogenous leukaemia (K562), triple-negative and ER-positive breast cancer (MDA-MB-321 and MCF7, respectively), colorectal carcinoma (DLD1, HCT116), cervical cancer (HeLa), and an embryonic kidney-derived transformed cell line (HEK293T)^41^. Cell line barcoding enabled the simultaneous identification and deconvolution of each cell line within the pooled sample, thereby eliminating inter-sample staining variability and enabling direct comparison under identical acquisition conditions (gating in Fig. S4B). Our results revealed substantial variation in the levels of individual marks across lines, with notable differences in H3K27me3, H3K9me3, H3K79me3, and H3K14ac, among others (Fig. 6M). Strikingly, multidimensional epigenetic space integration showed that cell lines derived from the same tissue of origin displayed markedly divergent epigenetic profiles (Fig. 6N). This is the case for the lung cancer lines (H460 and A594), colorectal lines (DLD1 and HCT116), and the two breast cancer lines (MDA-MB-321 and MCF7), which exhibited distinct epigenetic landscapes. This intra-cancer epigenetic heterogeneity likely reflects the genetic backgrounds, molecular subtypes, and mutational histories of each line. These results underscore the importance of epigenetic profiling as a complementary layer to transcriptomic or mutational classification in cancer research, which could be relevant to the clinical application of epigenetic drugs. Importantly, high-dimensional reduction showed that each cell line is separated in the UMAP space, with the sarcoma and HEK293T lines separated from the rest (Fig. 6O-P). This separation has biological meaning, as all epithelial and haematopoietic cancer lines cluster closer in the space, while the sarcoma line (U2OS), of mesenchymal origin, and the immortalised HEK293T line cluster separately from the rest^41^. In addition, unsupervised clustering identifies each cell line independently (Fig. 6Q), either in one or two clusters, thus suggesting that specific cancers could be identified by their epigenetic combinatorial signatures.

## DISCUSSION

Here, we present EpiFlow, a spectral flow cytometry-based platform for simultaneous single-cell quantification of 15 epigenetic markers, including histone modifications, DNA methylation and hydroxymethylation, and a total histone. Validation experiments confirmed that all antibodies in the panel can be deployed simultaneously without allosteric interference, that epigenetic marks detection is reliable across different chromatin compaction states and in response to epigenetic drugs, and that quantification can be achieved in whole-cell and nuclear preparations. In addition, we developed a multidimensional, integrative approach to visually assess changes in epigenetic landscapes using EpiFlow scores. Together, by leveraging the multiplexing capacity of spectral flow cytometry, EpiFlow offers a rapid, scalable, and quantitative approach to dissect epigenetic landscapes across diverse biological systems. We directly address the technological gap between low-throughput chromatin profiling methods and the growing demand for high-content, single-cell epigenetic analysis.

With the emergence of single-cell locus-specific methods such as scCUT&Tag and scATAC-seq, it is possible obtain genome-wide positional information for individual marks or chromatin accessibility. These are powerful tools for mapping regulatory elements at single-cell resolution, although limited to one or two marks per experiment, burdened by a complex library preparation process, and requiring substantial computational expertise for analysis. EpiFlow occupies a complementary and distinct niche: it sacrifices locus-level resolution in favour of the global and simultaneous quantification of 15 epigenetic marks, what constitutes a very powerful approach to study global epigenetic states, cellular heterogeneity, or drug-induced chromatin remodelling across large numbers of cells. EpiFlow offers advantages that sequencing-based methods cannot currently match, thanks to its straightforward and high-throughput acquisition, available to any laboratory with access to a spectral flow cytometer.

We have demonstrated that EpiFlow is a robust and versatile platform, substantially expanding the applicability of the previously described EpiTOF approach, which employed mass cytometry (CyTOF) to profile histone modifications^7^, and of earlier flow cytometry-based strategies, which were limited to a rather small number of histone marks^42^. While EpiTOF provided an important proof of concept for high-dimensional epigenetic profiling, mass cytometry remains costly due to the need for metal-conjugated antibodies and limited instrument accessibility, which restricts its widespread adoption. EpiFlow overcomes these barriers by leveraging conventional fluorophore-conjugated antibodies and spectral flow cytometers, which are increasingly available in research and clinical laboratories, thereby democratising multi-parametric single-cell epigenetic analysis. In addition, each of these previous methods were only tested in a single biological setting, either blood or ES cell differentiation. Importantly, EpiFlow is a versatile platform with room for the definition of specific, custom-designed panels, as shown here with two alternative EpiFlow panels. This versatility and epiprofiling capacity of EpiFlow is bound to increase in the near future as new and brighter fluorophores with improved spectral resolution continue to be developed. The EpiFlow platform can readily incorporate additional antibodies, further expanding the number of epigenetic marks that can be simultaneously interrogated in a single experiment.

The application of EpiFlow across diverse biological contexts demonstrates the platform’s capacity to deliver biological insight at a resolution that bulk methods cannot provide. In the epilepsy model, cell-type-resolved profiling revealed that glial and vascular cells, rather than neurons, are the primary sites of acute seizure-induced epigenetic reprogramming, a finding inaccessible to bulk tissue approaches. Similarly, the identification of tissue-specific epigenetic profiles in endothelial cells and resident macrophages across brain, lung, and liver demonstrates that even broadly defined cell types are epigenetically shaped by their tissue microenvironment^39,40^. These findings underscore the value of single-cell, multi-parametric epigenetic profiling as a discovery tool for cell-type-specific chromatin dynamics in physiology and disease.

As with any new technology, EpiFlow has limitations. EpiFlow requires cell fixation and permeabilisation, precluding functional or molecular analyses from the same cells; it cannot be combined with live-cell sorting for subsequent culture or sequencing without losing the epigenetic readout. Second, EpiFlow quantifies global levels of each mark rather than their genomic distribution, meaning that locus-specific changes that do not alter the total abundance of a modification will not be detected. And third, while we have demonstrated the absence of allosteric interference between antibodies in the panel, epitope competition or steric hindrance in specific chromatin contexts, for example, highly compacted heterochromatin in certain cell types, cannot be fully excluded and should be validated when EpiFlow is applied to new biological systems. These limitations should be considered when designing the further development of EpiFlow and highlight areas for future methodological refinement.

A particularly compelling application of EpiFlow lies in epigenetic drug discovery. We have provided proof-of-concept for the use of EpiFlow to detect on-target effects, off-target consequences, and dose- and time-dependent responses of epigenetic drugs with different specific or broad targets. The scalability of EpiFlow in a single experiment, its compatibility with standard multi-well plate formats, and its rapid acquisition times position it as a powerful tool for high-content epigenetic drug screening. In a field of intense pharmaceutical interest^43^, the ability to simultaneously monitor multiple chromatin modifications during drug treatment, capturing not only the intended effect but also secondary chromatin remodelling and off-target effects, could substantially accelerate the identification and characterisation of epigenetic modulators in both academic and industrial settings.

A key conceptual contribution of this work is the development of EpiFlow scores. By combining individual marks into biologically meaningful composite scores that reflect chromatin relaxation, heterochromatin, transcriptional states, and DNA methylation balance, we provide a layer of interpretation that goes beyond visualising individual marks. These scores reduce the complexity of a 15-dimensional epigenetic space into interpretable axes of chromatin biology, enabling rapid, intuitive comparison across conditions. Importantly, EpiFlow scores are sensitive to small yet synergistic changes that are missed when analysing marks individually, as demonstrated in the T2DM experiment (Fig. 4I). The scoring framework is modular and extensible: as new marks are incorporated into EpiFlow panels, additional or alternative scores can be defined to capture further dimensions of epigenetic regulation. We therefore propose that this integrative approach constitutes a general analytical framework for multi-parametric epigenetic data.

A central finding emerging from our heterogeneity experiments is that individual cell types can be defined by their epigenetic profiles alone, suggesting that specific combinations of histone modifications and DNA methylation states constitute cell-type-specific epigenetic codes or epigenetic cell identity. The divergent epigenetic landscapes observed across cancer cell lines derived from the same tissue of origin further support the notion that multi-parametric epigenetic profiling represents an independent classification layer, complementary to transcriptomic or mutational analyses. Notably, EpiFlow captured epigenetic states known to result from specific genomic or signalling alterations, such as the loss of H3K27me3 caused by PRC2 deletion in U2OS osteosarcoma cells^44^ or by AKT-mediated EZH2 inactivation downstream of PIK3CA mutation in H460 cells^45^, further demonstrating that multi-parametric epigenetic profiling can serve as a functional readout of underlying mutational landscapes. Beyond classification, the ability to define the specific epigenetic alterations that characterise individual cancers opens a route towards personalised therapeutic strategies: epigenetic changes identified at single-cell resolution could inform the selection of epigenetic drugs or combinations thereof in the context of resistant or refractory disease, for which chromatin dysregulation is increasingly recognised as both a driver and a target^2,43^.

Finally, the ability of EpiFlow to resolve cell-type-specific epigenetic states in accessible tissues such as peripheral blood opens significant translational opportunities. Blood-based profiling of immune cell subpopulations may capture both local immune dysregulation and systemic disease-associated chromatin alterations, offering a non-invasive diagnostic window. The prospect of defining a cell-type-specific epigenetic code in blood that reflects pathological states represents a compelling avenue for clinical biomarker development, particularly in conditions where tissue biopsies are impractical or where early detection is critical.

In summary, EpiFlow offers an unprecedented, high-throughput window into the single-cell epigenetic landscape. Its robustness across species and tissues, its scalability for drug screening, its capacity to dissect cellular heterogeneity, and its potential for biomarker discovery position it as a broadly applicable platform with impact spanning basic chromatin biology, pharmaceutical research, and translational medicine.

## METHODS

### Cell Culture

#### - Cell Lines

Jurkat T lymphoma cells (Jurkat) were maintained in RPMI media (Himedia, AT028), supplemented with 5% Fetal Bovine Serum (FBS; Gibco, A5256701) and 100 U/mL Penicillin (Panreac, A1837) and 100 μg/mL Streptomycin (P/S; Panreac, A1852). HeLa-S3 cells were maintained in DMEM (Himedia, AT066) media supplemented with 10% FBS and P/S. COS-7 cells were maintained in DMEM media supplemented with 5% FBS and P/S. MDCK cells were maintained in MEM (Gibco, 61100) media supplemented with 5% FBS, 2 mM glutamine 2 mM glutamine (TCI, G0063) and P/S. Neuro-2a (N2A), A549, HCT116, MCF7, MDA-MB-231, U2OS, HeLa, and HEK293 cells were maintained in DMEM media supplemented with 10% FBS, 2 mM glutamine and P/S. H460, MOLT-4, K562, and DLD-1 cells were maintained in RPMI media supplemented with 10% FBS, 2 mM glutamine and P/S. PC12 cells were maintained in RPMI media supplemented with 0.44 mM L-alanine (Merck, 1.010070), 0.4 mM L-asparagine (Merck, 1.015660), 0.4 mM L-proline (Merck, 1.074340), 0.4 mM L-aspartic acid (Panreac, A7219), 0.4 mM L-glutamic acid (Panreac, G8415), 2 mM glutamine, 5% Horse Serum (Gibco, 26050-088) and 5% FBS. Cells were grown in a humidified incubator containing 5% CO_2_ at 37°C and passaged when reaching 70-80% confluency. Hi5 insect cells were cultured in TC-100 medium (Lonza, 12-730Q) supplemented with 10% FBS in a shaker at 28° C.

#### - Yeast

*Saccharomyces cerevisiae* cells (W303-1a background) were grown in YPD medium (1% Bacto^TM^ yeast extract (Gibco, 212750), 2% Bacto^TM^ peptone (Gibco, 211677), and 2% glucose (Merck, 1.08342)) at 30° C in a shaking water bath.

#### - mESC

WT and histone H1 Triple Knock Out (H1TKO) E14 mouse Embryonic Stem Cells (mESC), cells derived from mouse strain 129/Ola blastocysts, were kindly provided by Arthur Skoultchi. Cells were grown in DMEM supplemented with 10% FBS, 0.44 mM L-alanine, 0.4 mM L-asparagine, 0.4 mM L-proline, 0.4 mM L-aspartic acid, 0.4 mM L-glutamic acid, 2 mM glutamine, 1 mM sodium pyruvate, 50 µM β-mercaptoethanol, P/S, and 103 U/mL ESGRO mLIF (Leukaemia Inhibitory Factor; Millipore, ESG1107) at 37° C and 5% CO_2_ on 0.1% gelatine-coated plates. Both cell types were grown until 75% of confluency.

### Arabidopsis thaliana growth

Arabidopsis thaliana Col-0 seeds were bleach-sterilized, stratified 2 days at 4°C and sown in agar plates containing 0.5x Murashige-Skoog medium (pH 5.7) supplemented with 0.5 g/L MES, 1% sucrose and 1% plant agar (Duchefa). Plants were grown vertically in a growth chamber at 21 °C and 60% moisture, under long-day conditions (16 h light/8 h dark cycles).

### Epigenetic drug treatments

Jurkat cells were treated with different epigenetic drugs for specific times. Cells were treated with 500 nM of Trichostatin A (TSA; Sigma-Aldrich, T8552; stock 5mM in DMSO) for 5 hours; 15 µM of Histone Lysine Demethylase Inhibitor VIII (GSK-J4; Merck, 420205; stock 200 mM in DMSO) for 24 hours; 15 µM of UNC0379 (Sigma-Aldrich, SML1465; stock 100 mM in DMSO) for 5 hours or with 1.46 µM, 7.3 µM and 36.5 µM for 5 and 8 hours; 300 nM of Cyclopentyl-(6,7-dichloro-4-pyridin-4-yl-phthalazin-1-yl)-amine (A-196; Sigma-Aldrich, SML1565; Stock 10 mM in DMSO) for 5 hours or with 28.8 nM, 144 nM and 720 nM for 5 and 8 hours. Aliquots were stored at −20°C upon reconstitution and protected from light until use.

### mESC prime-to-naïve de-differentiation

WT E14 mESC were maintained in primed-pluripotent state as described above. To obtain the naive-pluripotent state, 1 µM PD0325901 (MEK inhibitor; StemMACS Miltenyi Biotec, 130-103-923) and 3 µM CHIR99021 (GSK3 inhibitor; StemMACS Miltenyi Biotec, 130-103-926) were added to the medium. Medium was changed daily, and cells were split every 2 days. After 7 days of de-dedifferentiation cells were harvested.

### Animal Research

All procedures involving animals were conducted in accordance with European (2010/63/UE) and Spanish (RD 53/2013) legislation, in compliance with the ethical standards of Centro de Biología Molecular Severo Ochoa, and were performed under an authorised project licence (PROEX 328.1/23; PROEX 267.8/22; PROEX 014.1/24). C57BL/6J mice were kept in ventilated racks at 22±2°C and 55±10% humidity with a 12h/12h light cycle and free access to food and water at all times. The WT animals used in this project were euthanised by exposure to CO_₂_ and perfused intracardially with 0.9% NaCl. Organs were collected, snap frozen and stored at -80°C until use. Blood samples were also obtained and processed fresh.

#### - Type 2 diabetes mellitus

7-9-month-old C57BL/6J mice were subjected to a high-fat diet (HFD; 60% kcal from fat) for 14 weeks, at which time they received intraperitoneal injections of STZ (40□mg/kg) daily for over 5 consecutive days. After the STZ treatment, the animals were placed on an HFD for another 6 weeks. Animals were sacrificed at 12-14 months of age by exposure to CO_2_ and perfused intracardially with 0.9% NaCl. The liver was snap frozen and stored at -80° C until use. The animals used in this study were used in a previous study where we showed that they developed diabetes^46^.

#### - Immune humoral response

2-3-month-old C57BL/6J mice were immunised intraperitoneally with 2×10^9^ sheep red blood cells (SRBC; ThermoFisher, SR0051). Animals were sacrificed 7 days post-immunisation, and spleens were harvested and used fresh.

#### - Status epilepticus

4-month-old C57BL/6J mice were subjected to a behavioural assessment protocol designed to evaluate pentylenetetrazol (PTZ; Sigma-Aldrich, P6500)-induced seizure activity. The PTZ dose (60 mg/kg) and seizure scoring criteria were selected according to previously published methodology^47^.

Each animal was first recorded for 30 minutes prior to injection to establish an individual behavioural baseline in their home cages without any environmental enrichment. Following baseline acquisition, mice received an intraperitoneal injection of PTZ, a GABA_A receptor antagonist, dissolved in sterile 0.9% NaCl. Control animals received an intraperitoneal injection of 0.9% NaCl vehicle. Immediately after injection, animals were returned to their respective cages and recorded for 90 minutes. Seizure activity was scored according to a modified Racine scale, following the criteria described in the referenced study^47^. Behavioural analysis was performed by continuous observation of each individual mouse during the 10-minute analysis window. Latency to reach each Racine stage was recorded when applicable, as not all animals progressed through all stages. Racine stage 1 (immobility) was quantified as total duration of immobility. For Racine stages 2-6 (including head nodding, tail extension, rearing/kangaroo posture, and more severe generalized seizure manifestations), the number of discrete events was recorded. Racine stage 6 included mortality, consistent with published classifications. Animals were sacrificed 5 hours post-injection, perfused with 0.9% NaCl, and the brain was snap frozen and stored at -80° C until use.

### Antibody conjugation

Antibodies were conjugated to amino-reactive dyes (see Table S5) in 0.1 M sodium carbonate buffer (pH 8.3) for 1 hour at 30 molecules of dye to 1 molecule of antibody. Briefly, antibody buffers were exchanged for sodium carbonate buffer using Zeba 7K MWCO desalting columns (ThermoFisher, 89883), according to the manufacturer’s instructions. After conjugation, excess dye was removed and antibodies were brought back to the original storage buffer using Zeba 7K MWCO desalting columns. Antibody conjugation to Qdot705 (ThermoFisher, S10454), Biotin (Invitrogen, R10711), PE-Cy5 (Abcam, ab102893), PE-Cy5.5 (Abcam, ab102899) and PE-Cy7 (Abcam, ab102903) was performed according to the manufacturer’s instructions.

### Sample preparation for Flow cytometry

#### - Cell lines

Cells were trypsinised when necessary or directly pelleted if growing in suspension at 300 x g for 5 min. For the drug treatment experiments, cells were resuspended in PBS, incubated with viability dyes for 10 min at RT (see Table S5 for viability dye concentrations), and washed by centrifugation in PBS. Next, cells were fixed using Foxp3/Transcription Factor Fixation/Permeabilisation kit (Invitrogen, 00-5523- 00) for 10 min at RT and washed in FoxP3 wash buffer by centrifugation at 800 x g for 5 min. Veh- and TSA-treated Jurkat (Fig. 1) and mESC (Fig. 1,4) were subjected to epitope exposure by treating fixed cells with 2 N HCl for 10 min at RT and washed with FoxP3 wash buffer by centrifugation at 800 x g for 5 min. Cell concentration was determined in a Neubauer chamber, and 3.000 cells/µL were used for intracellular staining with EpiFlow antibodies, which was performed O/N on an orbital shaker (see Table S5 for antibodies and concentrations). The following morning, samples were washed in FoxP3 wash buffer and resuspended in FACS buffer (PBS containing 1% BSA and 1 mM EDTA) prior to data acquisition on an AURORA 5L.

#### - Arabidopsis thaliana nuclei isolation

Nuclei were isolated from 1 g of root tissue collected from 6-day-old *Arabidopsis thaliana*s eedlings using the CellLytic PN Isolation Kit (Sigma-Aldrich, CELLYTPN1). Tissue was finely chopped in 800 µL of 1× Nuclei Isolation Buffer (NIB) supplemented with 1 mM DL-Dithiothreitol (DTT; Sigma-Aldrich, D0632), then diluted with 4 mL NIB-DTT. The homogenate was processed in a KIMBLE Dounce tissue grinder (Sigma-Aldrich, D9063) with a loose pestle and filtered sequentially through 100 µm (Sysmex, 04-004-2328) and 30 µm (Sysmex, 04-004-2326) CellTicks filters. Triton X-100 (Sigma-Aldrich, T8787) was added to a final concentration of 0.1%. Next, nuclei were fixed in 1% methanol-free formaldehyde (Pierce, 28906) for 8 min at 4° C on a rocking platform and quenched with 0.125 M glycine. Subsequently, nuclei were pelleted by centrifugation at 1,960 x g for 30 min at 4 °C and resuspended in 2 mL of 2 N HCl for epitope exposer during 15min. Next, nuclei were washed with FACS buffer by centrifugation at 1,960 x g for 30 min at 4 °C. Nuclei concentration was determined by staining with DAPI and analysis in an AURORA 5L. Finally, 3.000 nuclei/µL were used for intracellular staining with EpiFlow antibodies, which was performed O/N on an orbital shaker (see Table S5 for antibodies and concentrations). The following morning, samples were washed in FoxP3 wash buffer and resuspended in FACS buffer prior to data acquisition on an AURORA 5L.

#### - Yeast

Exponentially growing cells, as estimated by microscopic observation, were collected by centrifugation at 1500 x g at RT for 3 min and subsequently washed with Milli-Q water. The number of cells was estimated using a counting chamber. 1 million cells pelleted and resuspended in Zymo-DTT buffer (10 mM NaCl, 10 mM Tris (pH 8), 1 mM EDTA, 40 mM DTT and 0.4µg/µL zymolyase 100T (MPBio, 083209-CF)) for yeast wall removal for 15 min. Next, cells were washed in PBS by centrifugation and fixed with FoxP3 fixation buffer for 10 min at RT. Subsequently, cells were washed with FoxP3 wash buffer by centrifugation at 3000 x g for 5 min at 4° C. Cells were next resuspended in 2 N HCl for epitope exposer during 15 min and washed with FoxP3 wash buffer by centrifugation at 3000 x g for 5 min at 4° C. Finally, cells were resuspended at 3.000 cells/µL in FoxP3 wash buffer and used for intracellular staining with EpiFlow antibodies, which was performed O/N on an orbital shaker (see Table S5 for antibodies and concentrations). The following morning, samples were washed in FoxP3 wash buffer and resuspended in FACS buffer prior to data acquisition on an AURORA 5L.

#### - Cell Cycle

For the experiment of the cell cycle, the HeLa cells at 70-80% of confluency were treated with 2 µM 5-Ethynyl-2′-deoxyuridine (EdU; Sigma-Aldrich, 900584) for 30 min. Then, cells were trypsinised and washed with cold PBS. The cells were fixed with 1% formaldehyde (FA; Sigma-Aldrich, 1.04003) and quenched at 0.125 M glycine. Cells were permeabilised with 0.25% Triton X-100 in PBS for 5 min, washed with Tris Buffer Saline (TBS), and incubated in 200 µl of Click-iT reaction (TBS, 2 mM CuSO4, 0.5 µM Alexa Fluor 488 azide (Jena Bioscience, CLK-1275-AZ-1) and 10 mM sodium ascorbate) for 30 minutes, at RT in the dark. Subsequently, samples were washed in PBS and incubated with 0.25 mg/ml RNase A (Qiagen, 1007885) in PBS at 37° C for 1 hour. Samples were washed in FACS buffer and cell concentration was determined in a Neubauer chamber. Finally, 3.000 cells/µL were used for intracellular staining with EpiFlow antibodies, which was performed O/N on an orbital shaker (see Table S5 for antibodies and concentrations). The following morning, samples were washed and resuspended in FACS buffer prior to data acquisition on an AURORA 5L.

#### - Splenic and Blood samples

For analysis of splenocyte populations, single-cell suspensions were prepared from homogenised spleens. For circulating cells, blood was collected in sodium heparin (Lab. Ramón Sala, 56.465) to prevent clotting. Cells were pelleted at 300 x g for 5 min and resuspended in erythrocyte lysis ACK buffer (0.15 M NH_4_Cl, 10 mM KHCO_3_, 0.1 mM EDTA; (pH 7.2-7.4)) and pelleted again. This step was performed twice. After obtaining splenocytes and circulating cells, samples were incubated with viability dyes for 10 min at room temperature in combination with a blocking antibody (see Table S5), and washed in FACS buffer by centrifugation. Samples were stained with surface antibodies for 15 min at 4° C in FACS buffer (see Table S5 for antibodies and concentrations). Antibodies were washed by centrifugation in FACS buffer (300 x g for 5 min), and cells were fixed using the Foxp3/Transcription Factor Fixation/Permeabilisation kit for 10 min at RT. After fixation, cells were washed and resuspended in FoxP3 wash buffer and cell concentration was determined in a Neubauer chamber. 3.000 splenocytes/µL or 15.000 circulating cells/µL were used for intracellular staining with EpiFlow antibodies, which was performed O/N on an orbital shaker (see Table S5 for antibodies and concentrations). The following morning, samples were washed in FoxP3 wash buffer and resuspended in FACS buffer prior to data acquisition on an AURORA 5L.

#### - Hepatocyte nuclei preparation

Nuclei isolation from frozen livers was adapted from Korenfeld et al. 2023^48^. Briefly, 1 mL of hypotonic buffer (250 mM sucrose, 10 mM Tris HCl (pH 7.8), 0.1% Igepal, 3 mM MgCl_2_, 10 mM KCl, 0.2 mM DTT, 0.5 mM spermidine and 1% (v/v) of protease inhibitor cocktail (ROCHE, 11836145001)) was added per 100 mg of tissue and homogenized in a KIMBLE Dounce tissue grinder set (15 strokes with loose pestle and 15 strokes with tight pestle). The homogenate was then passed through a 40 µm filter (Falcon™, 352340) (without pre-wetting), and the nuclei were collected by centrifugation at 500 x g for 10 min at 4°C. Nuclei pellets were resuspended by gentle pipetting in 2 mL of Nuclei Preparation Buffer+0.1%Tween (NPB; 10 mM Tris HCl (pH 7.5), 3 mM MgCl2, 10 mM NaCl, 0.1% Tween-20) per 100 mg of starting tissue, spun at 500 x g for 10 min at 4°C and resuspended in NPB Buffer. Nuclei were fixed by incubation in 1% PFA in PBS for 5 min, followed by an incubation for 5 min in 0.125 M glycine. The nuclei were washed twice in FACS buffer. Nuclei concentration was determined by staining with DAPI and analysis in an AURORA 5L. Finally, 3.000 nuclei/µL were used for intracellular staining with cell-type and EpiFlow antibodies, which was performed O/N on an orbital shaker (see Table S5 for antibodies and concentrations). The following morning, samples were washed in FACS buffer and resuspended in FACS buffer prior to data acquisition on an AURORA 5L.

#### - Tissue nuclei isolation

Total nuclei isolation from frozen brain, liver and lung was adapted from Nott et al. 2023^49^. Brain tissue was homogenised in lysis buffer (LB; 0.32 M sucrose, 5 mM CaCl_₂_, 3 mM magnesium acetate, 1 mM EDTA, 10 mM Tris-HCl (pH 8.0), 0.1% Triton X-100, 1% (v/v) of protease inhibitor cocktail and 3 mM DTT) using a wireless homogenising mortar. Liver and lung tissues were homogenised in LB by mechanical dissociation. The homogenate was then transferred to a KIMBLE Dounce tissue grinder set (15 strokes with loose pestle and 10 strokes with tight pestle). The nuclei were sequentially filtered through a 70 µm cell strainer (Falcon™, 35250) and a 40 µm cell strainer. The samples were then centrifuged at 800 × g for 10 minutes at 4° C. The nuclei pellet was washed three times with wash buffer (0.32 M sucrose, 5 mM CaCl_₂_, 3 mM magnesium acetate, 1 mM EDTA, 10 mM Tris-HCl (pH 8.0), and 1% (v/v) of protease inhibitor cocktail). Next, nuclei were fixed in 1% PFA in PBS for 10 minutes under gentle agitation. Fixation was quenched with 0.125 M glycine for 5 minutes. Fixed nuclei were centrifuged at 2.000 × g for 15 minutes at 4° C and washed once with FACS buffer by centrifugation at 2.000 × g for 15 minutes at 4° C. Nuclei concentration was determined by staining with DAPI and analysis in an AURORA 5L. Finally, 5.000 nuclei/µL from brain tissue and 3.000 nuclei/µL from liver and lung tissue were used for intracellular staining with cell-type and EpiFlow antibodies, which was performed O/N on an orbital shaker (see Table S5 for antibodies and concentrations). The following morning, samples were washed in FACS buffer and resuspended in FACS buffer prior to data acquisition on an AURORA 5L.

### Flow cytometry

All experiments were run on an AURORA 5L flow cytometer using SpectroFlow software (Cytek Biosciences). To optimise antibody concentration and obtain flow cytometry signals within the 10^4^-10^5^ intensity range, systematic titrations of all antibodies were performed from 1:50 to 1:10.000. For high-intensity signals, additional serial dilutions were performed.

#### - Data processing and analysis

Data were processed with EpiFlowProc-N/S (in-house software available upon request) and analysed using different approaches depending on the experimental context.

First, flow cytometry files (.fcs), with spillover matrices, were loaded into EpiFlowProc-N/S, an in-house, user-friendly program that generated new parameters, including histone PTM normalisations (histone PTM/total histone) and EpiFlow scores, based on compensated linear data at the single-event level. The software outputs a .csv file containing non-fluorescent parameters (SSCs and FSCs), compensated parameters, normalised parameters and/or EpiFlow scores. The resulting .csv files were subsequently converted back to .fcs format using FlowJo (BD Biosciences) or Floreada.io. FlowJo was used for cell population gating and geometric mean fluorescence intensity (MFI) calculation. Note that compensated parameters were evaluated using biexponential visualisation, whereas normalised data and EpiFlow Scores were analysed on a linear scale. Subsequent analysis steps were performed in Excel.

To compare epigenetic marks across samples with only one cell-type (i.e., cell lines), the MFI for each mark was first calculated for each sample. For each antibody, the MFI values were then summed across all cell population, and individual values were expressed as a percentage of this total, so that each row in the bubble plot sums to 100%. This normalisation removes differences in absolute MFI due to antibody performance or staining conditions, and allows comparison of the relative distribution of each histone mark across columns in a bubble plot.

For drug treatment experiments, MFI were obtained for each condition and expressed as a percentage of the vehicle. EpiFlow Scores were calculated in two steps. First, a grand mean was calculated using all replicates and conditions (vehicle and treated combined). Then each individual sample value was divided by this grand mean. This normalisation centres the data around the overall experimental average.

For complex samples containing multiple cell types, MFI were first corrected by the H2A+ population. This step accounts for differences in cell-type composition or antibody levels between samples. It also helps to exclude debris in nuclear preparations that are similar in size and complexity to nuclei. Importantly, H2A+ refers to a gated population based on total histone staining, which is gated prior to cell-type gating, not to total histone signal. The correction was performed separately for each antibody and each EpiFlow score. Specifically, each individual antibody signal or EpiFlow Score was divided by the corresponding H2A+ population value from the same sample. For experiments with a control or reference population, data were then expressed relative to that reference. This removes differences in absolute signal intensity between antibodies. For experiments without a control or reference population, the corrected signals from each cell type were summed within each sample and normalised to 100%. This allows comparison of the relative composition of cell types for each antibody. All EpiFlow Scores shown were calculated after H2A+ correction.

#### - High-dimensional reduction and clustering analysis

Data were processed using EpiFlowProc-Pro following a sequential pipeline including compensation, gating, asinh transformation, population definition and z-score scaling. Once they were scaled, the data were uploaded to OMIQ for high-dimensional reduction.

Using EpiFlowProc-Pro, the first step was using the *cell gating tab,* which manages data loading and gating. When .fcs files (experimental files and unstained controls) are uploaded, the software extracts spillover matrices and compensates linear data. Once compensated, population gating is performed based on non-fluorescent parameters (FSC and SSC parameters). In experiments including viability dyes, alive-cell selection can also be performed at this stage, using the same asinh transformation approach described below. Following gating, the *asinh transformation tab* applies an asinh transformation to samples using defined cofactors, allowing staining-specific transformations. Cofactors are determined by visual inspection of a histogram of the linear data for the selected population, enabling the user to define values for the entire sample set or for each individual sample. The histogram view also provides visual supervision of the transformed data. For cell-type markers (which display both negative and positive populations), a single histogram is shown. For epigenetic marks (which contain only a positive population), two histograms are displayed: one for the unstained control and one for the experimental sample. Once the transformation is complete, this tab defines negative and positive populations for each staining based on the selected cofactor. These populations are subsequently used in the next tab. In the *scaling tab*, z-score scaling is applied. By default, robust z-score scaling centred at 0 is used. The user may choose to centre either the negative or the positive population at a defined value. Typically, the negative population is centred for cell-type markers, whereas the positive population is centred for epigenetic marks. Importantly, z-score scaling preserves relative differences between individual cells, maintaining the within-sample distribution while adjusting the position of the data. This step is critical, as epigenetic mark signals depend on antibody efficiency and staining concentration, leading to differences in absolute signals levels, which may obscure the relative contribution of each mark, potentially biasing downstream dimensionality reduction and clustering analysis. Additional tabs in EpiFlowProc-Pro perform PTM normalisation and generate EpiFlow Score, as described for the EpiFlowProc-N/S version. Finally, the software exports a .csv file, allowing the user to select compensated, transformed, scaled, normalised, and/or score data for downstream analysis for the whole sample or for a gated population.

The exported .csv files were then uploaded into OMIQ (www.omiq.ai). Debris, dead cells, and aggregates were removed. Populations of interest were then manually gated, and high-dimensional reduction was performed using the UMAP algorithm, selecting only epigenetic marks. Unsupervised clustering was performed on the previously derived UMAPs parameters using ClusterX, together allowing visualisation of single-cell distribution in UMAP space via heat maps, and enabling comparison between manual gating of cell populations and unsupervised clustering by ClusterX.

### Histone PTM analysis by mass spectrometry

#### - Acidic protein extraction

Vehicle- and TSA-treated cells were trypsinized, collected by centrifuging at 300 x g for 5 min at RT, and washed twice with cold PBS by centrifugation. Next, cells were resuspended in hypotonic buffer (PBS, 20 mM Tris-HCl (pH 7.4), 10 mM NaCl, 3 mM MgCl_2_ and 1% (v/v) of protease inhibitor cocktail). After 15 min of incubation on ice, 25 µL of 10% Igepal (Sigma-Aldrich, I3021) was added, and the samples were mixed by vigorous vortexing for 10 seconds, then pelleted by centrifugation at 300 x g for 10 minutes at 4°C. Nuclei were washed once by centrifugation with hypotonic buffer. Next, 0.2 M H_2_SO_4_ was added at a 5:1 volumetric ratio and incubated in a wheel for 2 hours at 4°C. After acidic precipitation of proteins, samples were centrifuged at 3.400 x g for 10 min at 4°C. The supernatant was recovered, and basic proteins were precipitated with 20% trichloroacetic acid final concentration. The mixture was incubated O/N at 4° C. The following day, samples were centrifuged at 3.400 x g for 10 min at 4° C and the supernatant discarded. Next, the pellet was washed by resuspension in 100% acetone containing 0.1% HCl and centrifuged at 4.000 x g for 10 min at 4°C. Next, the pellet was washed twice with 100% acetone and air-dried after the second wash. Finally, samples were resuspended in 50 mM ammonium bicarbonate.

#### - Histone preparation for proteomics

Double derivatisation of lysine by propionic anhidride^50^ was performed on 20 µg of purified histones obtained by acid precipitation and resuspended in 100 mM ammonium bicarbonate (ABC). Samples were adjusted to 20 µL with ABC when necessary. Derivatisation was carried out using freshly prepared propionic anhydride:isopropanol (1:3, v/v). For each derivatisation step, 10 µL of the solution was added to the sample, followed by neutralisation with 25% ammonium hydroxide to maintain pH 8. Reactions were incubated for 15 min at 37°C with shaking (350 rpm) in a Thermomixer. Next, samples were dried down to 5 µL and resuspended in 20 µL of ABC. Derivatisation was repeated twice. Proteins were then digested overnight at 37 °C with 1 µg of trypsin in 100 mM ABC. Digestion was quenched with 2 µL of 25% trifluoroacetic acid, and samples were dried to <20 µL before being adjusted to 20 µL with ABC and pH 8. A second round of derivatisation was then performed using the same procedure (two consecutive derivatisation steps with propionic anhydride:isopropanol). Finally, samples were resuspended in 300 µL of 0.1% formic acid (buffer A), and 20 µL was loaded onto Evotips for subsequent liquid chromatography coupled to tandem mass spectrometry (LC-MS/MS) analysis.

#### - LC-MS/MS analysis

Samples were analysed using an Evosep One HPLC (Evosep) coupled to an Q Exactive HF Mass Spectrometer (Thermo Fisher Scientific) using an Endurance Evosep (EV1106) column (15 cm x 150 um ID) coupled to a stainless-steel emitter of 30 um ID and interfacecd with the Mass Spectrometer using Nanospray Flex Ion Source. Column was heated to maintain the temperature at 55°C. Peptides were eluted from Evotips and analysed using the Evosep One pre-programmed gradient for 30 samples per day (SPD). Data was acquired using a data-independent acquisition strategy. Full MS resolutions were set to 60,000 at m/z 200, and the full MS AGC target was 2e5 with a maximum injection time (IT) set to 200 ms. The AGC target value for fragment spectra was set to 3e6. 34 windows of 24 m/z scanning from 295 to 1115 m/z were employed with an overlap of 1 Da. MS2 resolution was set to 30,000, IT to 54 ms, and normalised collision energy (NCE) to 30%.

#### - Quantitative analysis

Data were further processed using Skyline 23.1.0.380 version (MacCoss lab, University of Washington, Seattle)^51^. The histone PTM Skyline template available at Panorama (https://panoramaweb.org/histone-ptm.url)^52,53^ was used to upload the raw files and extract the relevant histone PTM intensities. The histone marks of interest that were not already present in the template were added manually. Extracted Ion Chromatograms for the relevant marks were curated manually to contain unequivocally assigned fragments. MS2 intensities for the relevant peptides were exported for further processing into Excel. Quantification from the co-enriched iRT present in the sample and in the template was used to correct for unequal loading into the MS. Log2-normalised precursor intensities were used to quantify the TSA-induced changes in EpiFlow relevant marks. Briefly, log_₂_-transformed values were back-transformed by calculating 2^x^ to obtain linear intensity values for each peptide, which were then summed for each histone PTM of interest. For example, for lysine 4 of histone H3 (H3K4), we obtained summatory values for peptides containing H3K4un, H3K4ac, H3K4me1, H3K4me2, and H3K4me3. These sums were performed for each lysine of interest in vehicle and TSA-treated samples, and results were expressed as a percentage of the vehicle.

### Data Availability

The mass spectrometry proteomics data have been deposited to the ProteomeXchange Consortium via the PRIDE partner^54^ repository with the dataset identifier PXD075323.

### Statistical Analysis

Statistical analyses were performed using SPSS (IBM). Outliers were identified using the Explore tool (Tukey’s method). Homogeneity of variances was tested by the Levene test. Student’s t-test, assuming or not equal variances, was used to compare two groups. For more than two groups, one-way ANOVA was used for statistical analysis, followed by post-hoc comparisons assuming (Tukey’s) or not assuming equal variances (Games-Howell). All statistical analyses are two-tailed. In the figures, asterisks indicate p values as follows: ∗p□<□0.05; ∗∗p□<□0.01; ∗∗∗p□<□0.001.

## AUTHOR CONTRIBUTIONS

EP and ERB conceived this study. JRI and EP were involved in all experiments. LCM, JI, LM, JS, AGM, AA, JAT, CP, EL, MG and ERB were involved in cell line experiments, including mammals, insects, yeast, mES, cell cycle, and cancer studies. ERB, LCM and NMM were involved in B-cell and blood experiments. BD performed plant experiments. DGM and NRR were involved in brain/lung/liver experiments. ARM and JI were involved in T2DM experiments. VEZ and MDL were involved in epilepsy experiments. AMV, EC and JV performed proteomic experiments. EP, ERB, BRP and DJC contributed to the analysis design and EpiFlowProc development. EP wrote the manuscript, using artificial intelligence for English editing and with input from all authors.

## ACKNOWLEDGMENTS

We thank Elena Alejandre Ruíz (Palomer Lab), Raquel Nieto Pintado and Silvia Andrade Calvo (Flow Cytometry Core Facility at CBM), Laura Fernández Martín (Gómez Lab), and Prof Antonio Alcamí and Carolina Sánchez Fernandez (Alcamí Lab) for their support during this project. This project was funded by the Agencia Estatal de Investigación, Ministerio de Ciencia, Innovación y Universidades (MICIU) from the Spanish Government (MICIU/AEI/10.13039/501100011033): EP: PID2022-137384OA-I00 (ERDF /EU), RYC2021-031713-I (European Union NextGenerationEU/PRTR), EUR2024-153539 and CNS2024-154632; CP: PID2021-127824OB-I00 (ERDF/EU); MDL: PID2023- 149557OB-I00 (ERDF /EU); NMM: PID2024-162320OB-I00 (ERDF /EU) and CNS2022-135636 (European Union NextGenerationEU/PRTR); EL: PID2021-127824OB-I00 (ERDF /EU), CNS2022-135442 (European Union NextGenerationEU/PRTR) and PID2024-158067NB-I00 (ERDF /EU); NRR: PID2022-137552OA-100 (ERDF /EU); J.V: PID2021-122348NB-I00 (ERDF/EU), PID2024-155650NB-I00 (ERDF/EU) and EQC2021-007053-P; MG: PID2022-141380NB-I00 (ERDF/EU). Funded by Fundación Ramón Areces: EP: CIVP22A7609; NMM: CIVP22A7612. Funded by Comunidad de Madrid: J.V: S2022/BMD-7333-CM; AMV: “Ayudas de Atracción de Talento Investigador César Nombela 2023” (2023-T1/SAL-GL-28990). Funded by “la Caixa” Foundation: NMM: HR22-00447; J.V: LCF/PR/HR22/52420019. The CBM is supported by Fundación Ramón Areces and is a Severo Ochoa Center of Excellence (grant CEX2021-001154-S funded by MICIU/AEI/10.13039/501100011033). The CNIC is supported by the Instituto de Salud Carlos III (ISCIII), the Ministerio de Ciencia, Innovación Y Universidades (MICIU) and the Pro CNIC Foundation), and is a Severo Ochoa Center of Excellence (grant CEX2020-001041-S funded by MICIU/AEI/10.13039/501100011033).

## Notes

### Competing Interest Statement

The authors have declared no competing interest.

## REFERENCES

1. Allis, C.D., and Jenuwein, T. (2016). The molecular hallmarks of epigenetic control. Nature Reviews Genetics 2016 17:8 17, 487–500. 10.1038/nrg.2016.59.

2. Laisné, M., Lupien, M., and Vallot, C. (2024). Epigenomic heterogeneity as a source of tumour evolution. Nature Reviews Cancer 2024 25:1 25, 7–26. 10.1038/s41568-024-00757-9.

3. Lu, C., Coradin, M., Porter, E.G., and Garcia, B.A. (2021). Accelerating the Field of Epigenetic Histone Modification through Mass Spectrometry–Based Approaches. Molecular and Cellular Proteomics 20, 100006. 10.1074/MCP.R120.002257/ASSET/832353A1-1571-41D3-A55E-4C6D0C1DB00E/MAIN.ASSETS/GR4.JPG.

4. Kaya-Okur, H.S., Wu, S.J., Codomo, C.A., Pledger, E.S., Bryson, T.D., Henikoff, J.G., Ahmad, K., and Henikoff, S. (2019). CUT&Tag for efficient epigenomic profiling of small samples and single cells. Nature Communications 2019 10:1 10, 1930-. 10.1038/s41467-019-09982-5.

5. Mehrmohamadi, M., Sepehri, M.H., Nazer, N., and Norouzi, M.R. (2021). A Comparative Overview of Epigenomic Profiling Methods. Front. Cell Dev. Biol. 9, 714687. 10.3389/FCELL.2021.714687/TEXT.

6. Orsburn, B.C. (2025). Simultaneous single-cell proteomics and epigenetic analysis of histone deacetylase inhibition in human cells. Communications Biology 2025 9:1 9, 108-. 10.1038/s42003-025-09452-3.

7. Cheung, P., Vallania, F., Warsinske, H.C., Donato, M., Schaffert, S., Chang, S.E., Dvorak, M., Dekker, C.L., Davis, M.M., Utz, P.J., et al. (2018). Single-Cell Chromatin Modification Profiling Reveals Increased Epigenetic Variations with Aging. Cell 173, 1385–1397.e14. 10.1016/J.CELL.2018.03.079/ATTACHMENT/67CCE4FC-B351-4A36-A467-706CB7B678F8/MMC5.PDF.

8. Sharma, S., Boyer, J., and Teyton, L. (2023). A practitioner’s view of spectral flow cytometry. Nature Methods 2023 21:5 21, 740–743. 10.1038/s41592-023-02042-3.

9. Bannister, A.J., and Kouzarides, T. (2011). Regulation of chromatin by histone modifications. Cell Research 2011 21:3 21, 381–395. 10.1038/cr.2011.22.

10. Yoshida, M., Kijima, M., Akita, M., and Beppu, T. (1990). Potent and specific inhibition of mammalian histone deacetylase both in vivo and in vitro by trichostatin A. Journal of Biological Chemistry 265, 17174–17179. 10.1016/S0021-9258(17)44885-X.

11. Fan, Y., Nikitina, T., Zhao, J., Fleury, T.J., Bhattacharyya, R., Bouhassira, E.E., Stein, A., Woodcock, C.L., and Skoultchi, A.I. (2005). Histone H1 Depletion in Mammals Alters Global Chromatin Structure but Causes Specific Changes in Gene Regulation. Cell 123, 1199–1212. 10.1016/J.CELL.2005.10.028.

12. Moshi, J.M., Ummelen, M., Broers, J.L.V., Ramaekers, F.C.S., and Hopman, A.H.N. (2023). Impact of antigen retrieval protocols on the immunohistochemical detection of epigenetic DNA modifications. Histochem. Cell Biol. 159, 513–526. 10.1007/S00418-023-02187-4.

13. Capuano, F., Mülleder, M., Kok, R., Blom, H.J., and Ralser, M. (2014). Cytosine DNA methylation is found in Drosophila melanogaster but absent in Saccharomyces cerevisiae, Schizosaccharomyces pombe, and other yeast species. Anal. Chem. 86, 3697–3702. 10.1021/AC500447W.

14. Erdmann, R.M., Souza, A.L., Clish, C.B., and Gehring, M. (2015). 5-hydroxymethylcytosine is not present in appreciable quantities in Arabidopsis DNA. G3: Genes, Genomes, Genetics 5, 1–8. 10.1534/G3.114.014670/-/DC1.

15. Fu, Y., Yang, Y., Zhang, H., Farley, G., Wang, J., Quarles, K.A., Weng, Z., and Zamore, P.D. (2018). The genome of the Hi5 germ cell line from trichoplusia ni, an agricultural pest and novel model for small RNA biology. Elife 7. 10.7554/ELIFE.31628.

16. Heinemann, B., Nielsen, J.M., Hudlebusch, H.R., Lees, M.J., Larsen, D.V., Boesen, T., Labelle, M., Gerlach, L.O., Birk, P., and Helin, K. (2014). Inhibition of demethylases by GSK-J1/J4. Nature 2014 514:7520 514, E1–E2. 10.1038/nature13688.

17. Ma, A., Yu, W., Li, F., Bleich, R.M., Herold, J.M., Butler, K. V., Norris, J.L., Korboukh, V., Tripathy, A., Janzen, W.P., et al. (2014). Discovery of a selective, substrate-competitive inhibitor of the lysine methyltransferase SETD8. J. Med. Chem. 57, 6822–6833. 10.1021/JM500871S/SUPPL_FILE/JM500871S_SI_001.PDF.

18. Bromberg, K.D., Mitchell, T.R.H., Upadhyay, A.K., Jakob, C.G., Jhala, M.A., Comess, K.M., Lasko, L.M., Li, C., Tuzon, C.T., Dai, Y., et al. (2017). The SUV4-20 inhibitor A-196 verifies a role for epigenetics in genomic integrity. Nature Chemical Biology 2017 13:3 13, 317–324. 10.1038/nchembio.2282.

19. Serrano, L., Martínez-Redondo, P., Marazuela-Duque, A., Vazquez, B.N., Dooley, S.J., Voigt, P., Beck, D.B., Kane-Goldsmith, N., Tong, Q., Rabanal, R.M., et al. (2013). The tumor suppressor SirT2 regulates cell cycle progression and genome stability by modulating the mitotic deposition of H4K20 methylation. Genes Dev. 27, 639–653. 10.1101/gad.211342.112.

20. Alabert, C., and Groth, A. (2012). Chromatin replication and epigenome maintenance. Nature Reviews Molecular Cell Biology 2012 13:3 13, 153–167. 10.1038/nrm3288.

21. Greenberg, M.V.C., and Bourc’his, D. (2019). The diverse roles of DNA methylation in mammalian development and disease. Nature Reviews Molecular Cell Biology 2019 20:10 20, 590–607. 10.1038/s41580-019-0159-6.

22. Bachman, M., Uribe-Lewis, S., Yang, X., Williams, M., Murrell, A., and Balasubramanian, S. (2014). 5-Hydroxymethylcytosine is a predominantly stable DNA modification. Nature Chemistry 2014 6:12 6, 1049–1055. 10.1038/nchem.2064.

23. Bannister, A.J., and Kouzarides, T. (2011). Regulation of chromatin by histone modifications. Cell Research 2011 21:3 21, 381–395. 10.1038/cr.2011.22.

24. Reverón-Gómez, N., González-Aguilera, C., Stewart-Morgan, K.R., Petryk, N., Flury, V., Graziano, S., Johansen, J.V., Jakobsen, J.S., Alabert, C., and Groth, A. (2018). Accurate Recycling of Parental Histones Reproduces the Histone Modification Landscape during DNA Replication. Mol. Cell 72, 239–249.e5. 10.1016/J.MOLCEL.2018.08.010/ATTACHMENT/613A9079-956F-4997-8CC8-189D88786930/MMC2.PDF.

25. Theunissen, T.W., Friedli, M., He, Y., Planet, E., O’Neil, R.C., Markoulaki, S., Pontis, J., Wang, H., Iouranova, A., Imbeault, M., et al. (2016). Molecular Criteria for Defining the Naive Human Pluripotent State. Cell Stem Cell 19, 502–515. 10.1016/j.stem.2016.06.011.

26. Tosolini, M., Brochard, V., Adenot, P., Chebrout, M., Grillo, G., Navia, V., Beaujean, N., Francastel, C., Bonnet-Garnier, A., and Jouneau, A. (2018). Contrasting epigenetic states of heterochromatin in the different types of mouse pluripotent stem cells. Scientific Reports 2018 8:1 8, 5776-. 10.1038/s41598-018-23822-4.

27. Novo, C.L., Tang, C., Ahmed, K., Djuric, U., Fussner, E., Mullin, N.P., Morgan, N.P., Hayre, J., Sienerth, A.R., Elderkin, S., et al. (2016). The pluripotency factor Nanog regulates pericentromeric heterochromatin organization in mouse embryonic stem cells. Genes Dev. 30, 1101–1115. 10.1101/GAD.275685.115.

28. Béguelin, W., Popovic, R., Teater, M., Jiang, Y., Bunting, K.L., Rosen, M., Shen, H., Yang, S.N., Wang, L., Ezponda, T., et al. (2013). EZH2 Is Required for Germinal Center Formation and Somatic EZH2 Mutations Promote Lymphoid Transformation. Cancer Cell 23, 677–692. 10.1016/j.ccr.2013.04.011.

29. Jiang, Y., Ortega-Molina, A., Geng, H., Ying, H.Y., Hatzi, K., Parsa, S., McNally, D., Wang, L., Doane, A.S., Agirre, X., et al. (2017). CREBBP inactivation promotes the development of HDAC3-dependent lymphomas. Cancer Discov. 7, 38–53. 10.1158/2159-8290.CD-16-0975/333294/AM/CREBBP-INACTIVATION-PROMOTES-THE-DEVELOPMENT-OF.

30. Kulis, M., Merkel, A., Heath, S., Queirós, A.C., Schuyler, R.P., Castellano, G., Beekman, R., Raineri, E., Esteve, A., Clot, G., et al. (2015). Whole-genome fingerprint of the DNA methylome during human B cell differentiation. Nature Genetics 2015 47:7 47, 746–756. 10.1038/ng.3291.

31. Lai, A.Y., Mav, D., Shah, R., Grimm, S.A., Phadke, D., Hatzi, K., Melnick, A., Geigerman, C., Sobol, S.E., Jaye, D.L., et al. (2013). DNA methylation profiling in human B cells reveals immune regulatory elements and epigenetic plasticity at Alu elements during B-cell activation. Genome Res. 23, 2030–2041. 10.1101/GR.155473.113.

32. Price, M.J., Scharer, C.D., Kania, A.K., Randall, T.D., and Boss, J.M. (2021). Conserved epigenetic programming and enhanced heme metabolism drive memory B cell reactivation. J. Immunol. 206, 1493. 10.4049/JIMMUNOL.2000551.

33. Scharer, C.D., Barwick, B.G., Guo, M., Bally, A.P.R., and Boss, J.M. (2018). Plasma cell differentiation is controlled by multiple cell division-coupled epigenetic programs. Nature Communications 2018 9:1 9, 1698-. 10.1038/s41467-018-04125-8.

34. Barwick, B.G., Scharer, C.D., Martinez, R.J., Price, M.J., Wein, A.N., Haines, R.R., Bally, A.P.R., Kohlmeier, J.E., and Boss, J.M. (2018). B cell activation and plasma cell differentiation are inhibited by de novo DNA methylation. Nature Communications 2018 9:1 9, 1900-. 10.1038/s41467-018-04234-4.

35. Zhao, Q., Cui, X., Zhu, Q., Li, F., Bao, R., Shi, T., Liu, H., Lv, W., Xu, Y., Gao, Y., et al. (2025). Non-catalytic mechanisms of KMT5C regulating hepatic gluconeogenesis. Nature Communications 2025 16:1 16, 1483-. 10.1038/s41467-025-56696-y.

36. Ling, C., and Rönn, T. (2019). Epigenetics in Human Obesity and Type 2 Diabetes. Cell Metab. 29, 1028–1044. 10.1016/J.CMET.2019.03.009.

37. Hauser, R.M., Henshall, D.C., and Lubin, F.D. (2018). The Epigenetics of Epilepsy and Its Progression. Neuroscientist 24, 186–200. 10.1177/1073858417705840/ASSET/9BA74D16-D9A7-4074-BF3D-67BB7AFD3A6D/ASSETS/IMAGES/LARGE/10.1177_1073858417705840-FIG2.JPG.

38. Kobow, K., Kaspi, A., Harikrishnan, K.N., Kiese, K., Ziemann, M., Khurana, I., Fritzsche, I., Hauke, J., Hahnen, E., Coras, R., et al. (2013). Deep sequencing reveals increased DNA methylation in chronic rat epilepsy. Acta Neuropathol. 126, 741–756. 10.1007/S00401-013-1168-8/FIGURES/6.

39. Paik, D.T., Tian, L., Williams, I.M., Rhee, S., Zhang, H., Liu, C., Mishra, R., Wu, S.M., Red-Horse, K., and Wu, J.C. (2020). Single-Cell RNA Sequencing Unveils Unique Transcriptomic Signatures of Organ-Specific Endothelial Cells. Circulation 142, 1848–1862. 10.1161/CIRCULATIONAHA.119.041433/SUPPL_FILE/CIRC_CIRCULATIONAHA-2019-041433_SUPP4.XLSX.

40. Lavin, Y., Winter, D., Blecher-Gonen, R., David, E., Keren-Shaul, H., Merad, M., Jung, S., and Amit, I. (2014). Tissue-resident macrophage enhancer landscapes are shaped by the local microenvironment. Cell 159, 1312–1326. 10.1016/J.CELL.2014.11.018/ATTACHMENT/C809A61F-F547-49B3-A1D1-34D4FE4F2EA4/MMC9.PDF.

41. Barretina, J., Caponigro, G., Stransky, N., Venkatesan, K., Margolin, A.A., Kim, S., Wilson, C.J., Lehár, J., Kryukov, G. V., Sonkin, D., et al. (2012). The Cancer Cell Line Encyclopedia enables predictive modelling of anticancer drug sensitivity. Nature 2012 483:7391 483, 603–607. 10.1038/nature11003.

42. Golden, C.S., Williams, S., Sreerama, S., Blankevoort, S., Yost, H.J., Tristani-Firouzi, M., Belkina, A., and Serrano, M.A. (2026). Nuclear Histone 3 Post-Translational Modification Profiling in Whole Cells using Spectral Flow Cytometry. bioRxiv, 2024.10.03.616268. 10.1101/2024.10.03.616268.

43. Dai, W., Qiao, X., Fang, Y., Guo, R., Bai, P., Liu, S., Li, T., Jiang, Y., Wei, S., Na, Z., et al. (2024). Epigenetics-targeted drugs: current paradigms and future challenges. Signal Transduction and Targeted Therapy 2024 9:1 9, 332-. 10.1038/s41392-024-02039-0.

44. Feng, H., Tillman, H., Wu, G., Davidoff, A.M., Yang, J., Feng, H., Tillman, H., Wu, G., Davidoff, A.M., and Yang, J. (2018). Frequent epigenetic alterations in polycomb repressive complex 2 in osteosarcoma cell lines. Oncotarget 9, 27087–27091. 10.18632/ONCOTARGET.25484.

45. Cha, T.L., Zhou, B.P., Xia, W., Wu, Y., Yang, C.C., Chen, C. Te, Ping, B., Otte, A.P., and Hung, M.C. (2005). Molecular biology: Akt-mediated phosphorylation of EZH2 suppresses methylation of lysine 27 in histone H3. Science (1979). 310, 306–310. 10.1126/SCIENCE.1118947/SUPPL_FILE/CHA.SOM.PDF.

46. Carús-Cadavieco, M., González de la Fuente, S., Berenguer López, I., Serrano-Lope, M.A., Aguado, B., Guix, F., Palomer, E., and Dotti, C.G. (2024). Loss of Cldn5 -and increase in Irf7-in the hippocampus and cerebral cortex of diabetic mice at the early symptomatic stage. Nutrition & Diabetes 2024 14:1 14, 64-. 10.1038/s41387-024-00325-y.

47. Van Erum, J., Van Dam, D., and De Deyn, P.P. (2019). PTZ-induced seizures in mice require a revised Racine scale. Epilepsy & Behavior 95, 51–55. 10.1016/J.YEBEH.2019.02.029.

48. Korenfeld, N., Toft, N.I., Dam, T. V., Charni-Natan, M., Grøntved, L., and Goldstein, I. (2023). Protocol for bulk and single-nuclei chromatin accessibility quantification in mouse liver tissue. STAR Protoc. 4, 102462. 10.1016/J.XPRO.2023.102462.

49. Nott, A., Schlachetzki, J.C.M., Fixsen, B.R., and Glass, C.K. (2021). Nuclei isolation of multiple brain cell types for omics interrogation. Nature Protocols 2021 16:3 16, 1629–1646. 10.1038/s41596-020-00472-3.

50. Karch, K.R., Sidoli, S., and Garcia, B.A. (2016). Identification and Quantification of Histone PTMs Using High-Resolution Mass Spectrometry. Methods Enzymol. 574, 3–29. 10.1016/BS.MIE.2015.12.007.

51. Pino, L.K., Searle, B.C., Bollinger, J.G., Nunn, B., MacLean, B., and MacCoss, M.J. (2020). The Skyline ecosystem: Informatics for quantitative mass spectrometry proteomics. Mass Spectrom. Rev. 39, 229–244. 10.1002/MAS.21540.

52. Sharma, V., Eckels, J., Schilling, B., Ludwig, C., Jaffe, J.D., MacCoss, M.J., and MacLean, B. (2018). Panorama Public: A Public Repository for Quantitative Data Sets Processed in Skyline. Molecular & Cellular Proteomics 17, 1239–1244. 10.1074/MCP.RA117.000543.

53. Panorama Dashboard: /Panorama Public/2024/U of Pennsylvania Epigenetics Institute - Histone PTM Skyline template https://panoramaweb.org/Panorama%20Public/2024/U%20of%20Pennsylvania%20Epigenetics%20Institute%20-%20Histone%20PTM%20Skyline%20template/project-begin.view.

54. Perez-Riverol, Y., Bandla, C., Kundu, D.J., Kamatchinathan, S., Bai, J., Hewapathirana, S., John, N.S., Prakash, A., Walzer, M., Wang, S., et al. (2025). The PRIDE database at 20 years: 2025 update. Nucleic Acids Res. 53, D543–D553. 10.1093/NAR/GKAE1011.

