## Supplementary Figures for "EpiFlow: multidimensional single-cell epigenetic profiling by spectral flow cytometry"

### Supplementary Figure S1

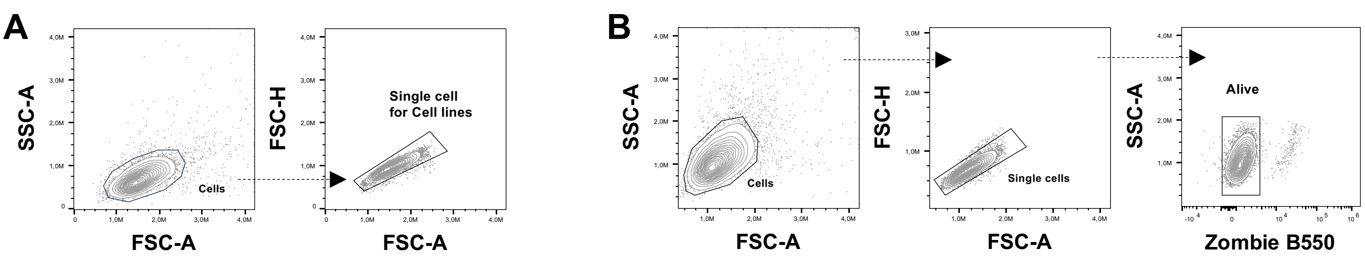


**C**

```
Human  (Homo sapiens)                MARTKQTARKSTGGKAPRKQLATKAARKSAPATGGVKKPHRYRPGTVALREIRRYQKSTELLIRKLPFQRLVREIAQDFKT 80
Monkey (Chlorocebus sabaeus )        MARTKQTARKSTGGKAPRKQLATKAARKSAPSTGGVKKPHRYRPGTVALREIRRYQQSTELLICKLPFQRLVQEIAQDFKT 80
Dog  (Canis lupus familiaris )       MARTKQTAHKSTGGKAPRKQLATKAARKSAPSTGGVKKPHRYRPGTVALHEIRHYQKSTELLIRKFPFQRLVREIAQDFKT 80
Rat  (Rattus norvegicus )            MARTKQTARKSTGGKAPRKQLATKAARKSAPATGGVKKPHRYRPGTVALREIRRYQKSTELLIRKLPFQRLVREIAQDFKT 80
Mouse  (Mus musculus )               MARTKQTARKSTGGKAPRKQLATKAARKSAPATGGVKKPHRYRPGTVALREIRRYQKSTELLIRKLPFQRLVREIAQDFKT 80
Insect (Trichoplusia ni)             MARTKQTARKSTGGKAPRKQLATKAARKSAPSTGGVKKPHRYRPGTVALREIRRYQKSTELLIRKLPFQRLVREIAQDFKT 80
Plant  (Arabidopsis thaliana)        MARTKQTARKSTGGKAPRKQLATKAARKSAPATGGVKKPHRFRPGTVALREIRKYQKSTELLIRKLPFQRLVREIAQDFKT 80
Yeast  (S. cerevisiae)               MARTKQTARKSTGGKAPRKQLASKAARKSAPSTGGVKKPHRYKPGTVALREIRRFQKSTELLIRKLPFQRLVREIAQDFKT 80
                                     ********:*************:********:*********::******:***::*:****** *:******:******** 80
```

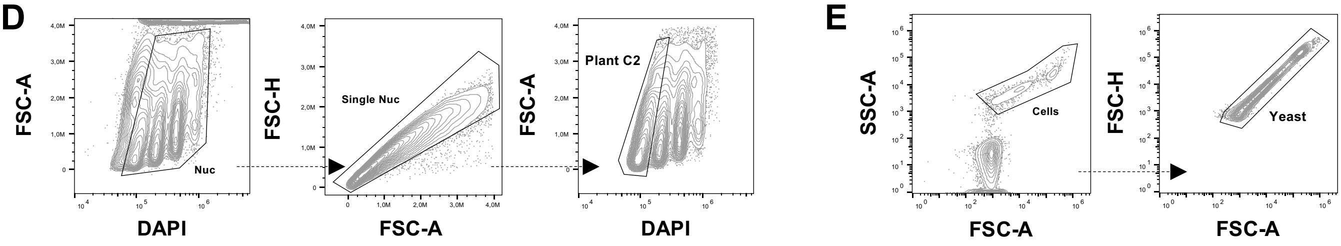


**F**

**Chromatin Relaxation (CR):**

$$\frac{H3K9ac + H3K14ac + H3K27ac + H4K8ac + H4K16ac}{H3K9me3 + H3K27me3 + H4K20me3}$$

**DNA methylation (DNA Meth):**

$$\frac{5\text{-}hmC}{5\text{-}mC}$$

**Facultative Heterochromatin Activation (FHcA):**

$$\frac{H3K27ac}{H3K27me3}$$

**Heterochromatin Index (HcI):**

$$H3K9me3 + H3K27me3 + H4K20me3 + 5mC$$

**Enhancer Activity (EA):**

$$\frac{H3K27ac}{H3K4me3}$$

**Facultative vs Constitutive Heterochromatin (FHc/CHc):**

$$\frac{H3K27me3}{H3K9me3}$$

**Initiation & Elongation index (I&E):**

$$H3K4me3 + H3K36me2 + H3K36me3 + H3K79me3 + 5hmC$$

**Transcriptional Potential (TP):**

$$\frac{H3K4me3 + H3K9ac + H3K14ac + H3K27ac + H3K36me2 + H3K36me3 + H3K79me3 + H4K8ac + H4K16ac}{H3K9me3 + H3K27me3 + H4K20me3}$$

**Supplementary Figure S1: Cross-species histone conservation, EpiFlow scores and gating strategies. A)** Representative gating strategies of diverse cell types, including Jurkat, COS7, MDCK, PC12, N2A and Hi5. **B)** Representative plots showing the gating strategies of cell lines treated with epigenetic drugs. **C)** ClustalW alignment of histone H3.1 across eukaryotes. **D-E)** Representative plots showing gating strategies for plants (D) and yeast (E). **F)** Representation of the histone marks and DNA methylations used for calculating each EpiFlow Score.

### Supplementary Figure S2

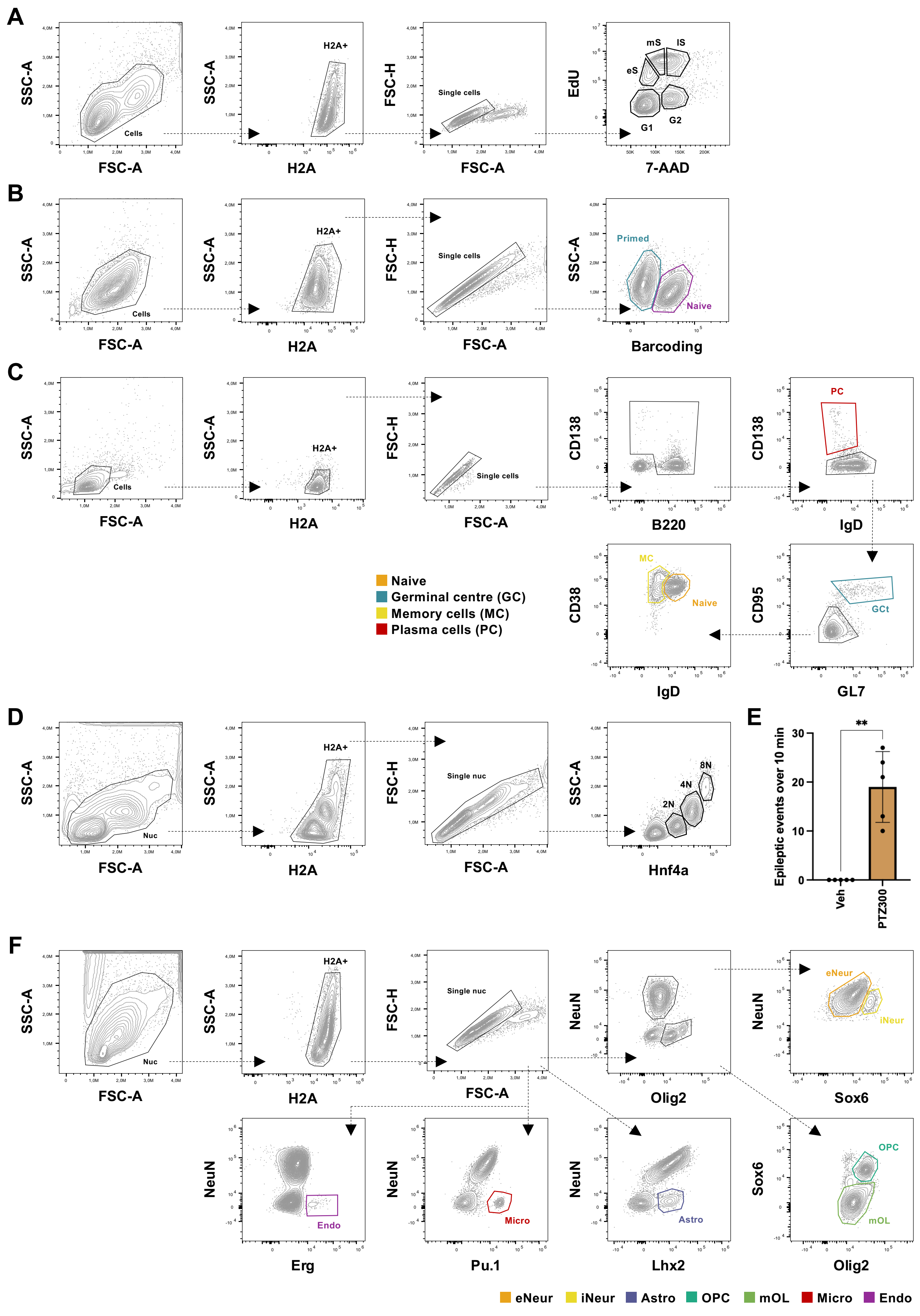

**Supplementary Figure S2: Gating strategies for relevant biomedical research settings. A-D)** Representative gating strategy plots for HeLa-S3 used in cell cycle experiments (A) to identify G1, early S (eS), middle S (mS), late S (lS), and G2 phases; in mouse ES cells (B) to identify primed (cyan) and naive (magenta); in splenic samples (C) to identify naive (orange), germinal centre (GC, cyan), memory cells (MC, yellow) and plasma cells (PC, magenta); in liver samples (D) to identify hepatocytes 2N, 4N, and 8N. **E)** Bar plot showing the number of epileptic seizures in vehicle (Veh) and PTZ-treated animals over the first 10 minutes post-PTZ administration (N indicated in the plot). Statistical analysis was performed using Student's t-test, **$p < 0.01$. **F)** Representative gating strategy plots used in brain samples (E) to identify excitatory neurons (eNeur, orange), inhibitory neurons (iNeur, yellow), astrocytes (Astro, purple), oligodendrocyte precursor cells (OPC, turquoise green), mature oligodendrocytes (mOL, pale green), microglia (Micro, red) and endothelial cells (Endo, magenta).

#### Supplementary Figure S3

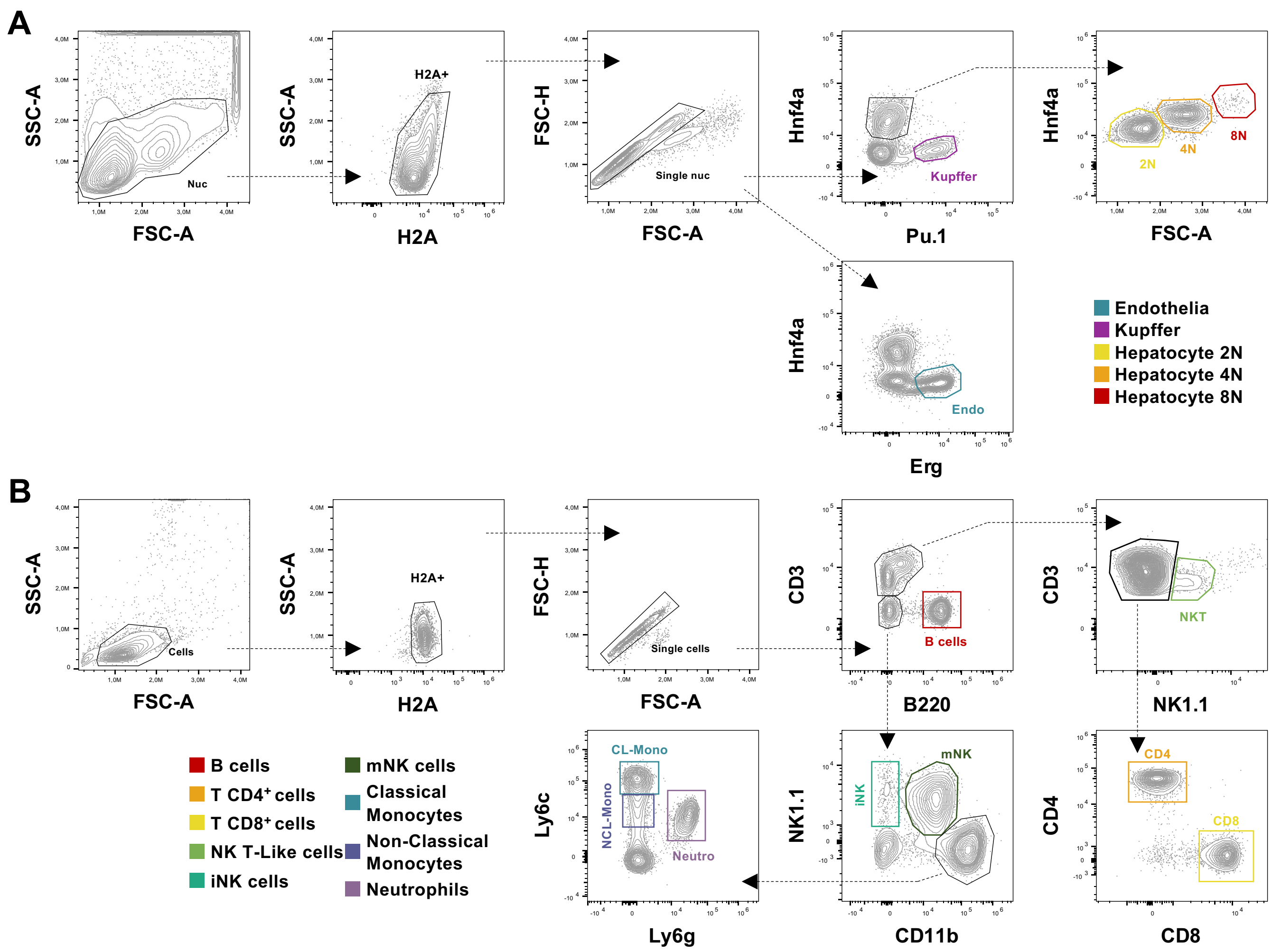


**Supplementary Figure S3: Gating strategies for cell heterogeneity in naturally occurring complex samples.** Representative gating strategy plots used in liver (A) to identify endothelial cells (cyan), Kupffer cells (magenta), hepatocytes 2N (yellow), 4N (orange) and 8N (red), and in blood (B) to identify B cells (red), CD4 T cells (orange), CD8 T cells (yellow), natural killer (NK) T-like cells (pale green), immature NK cells (iNK, turquoise green), mature NK cells (mNK, dark green), classical monocytes (CL-Mono, blue), non-classical monocytes (NCL-Mono, dark purple), and neutrophils (Neutro, pale purple).

### Supplementary Figure S4

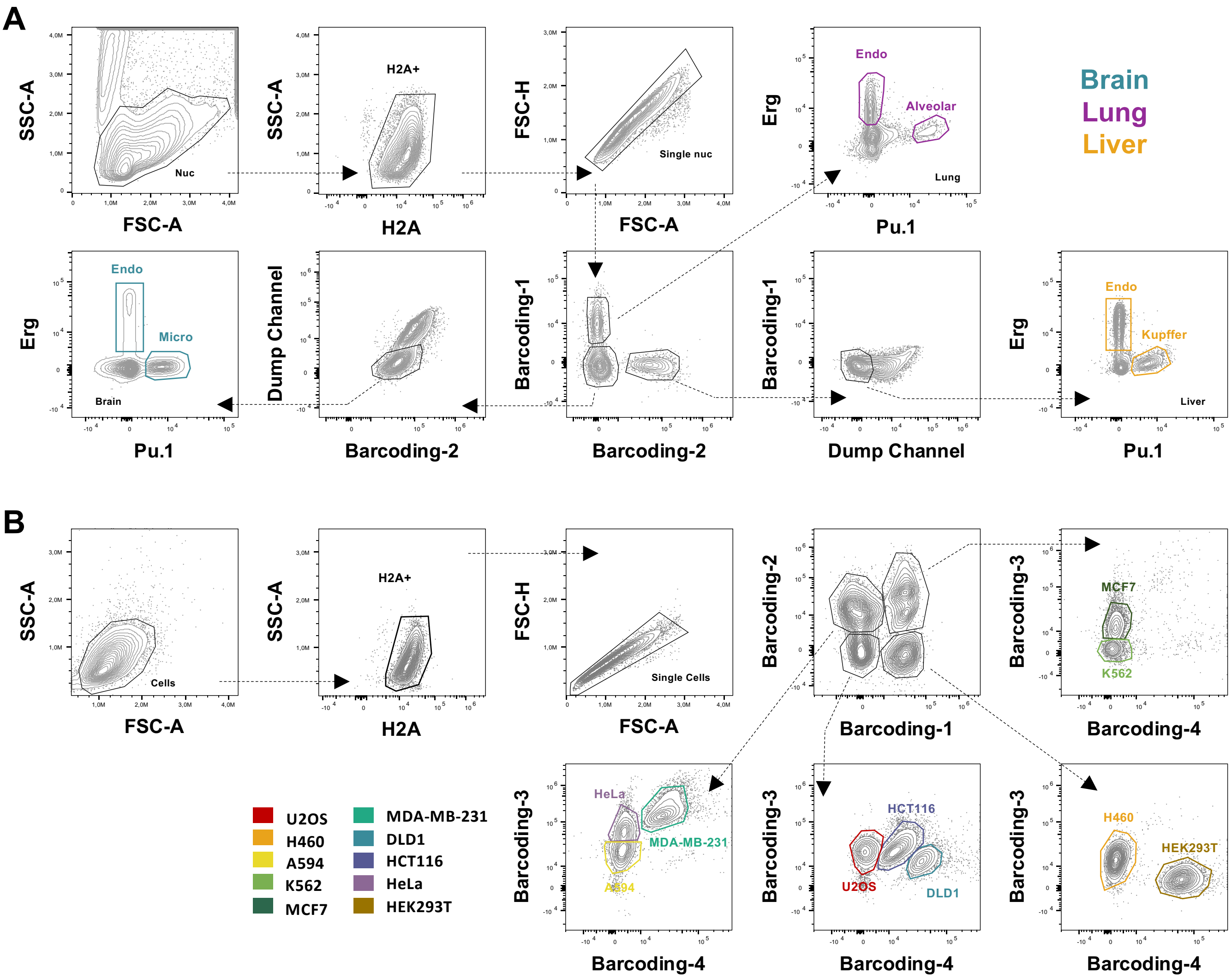


**Supplementary Figure S4: Gating strategies for artificial mixtures.** Representative gating strategy plots and barcoding deconvolution used to identify tissue-specific endothelial cells and tissue resident macrophages (A) in the mouse brain (cyan), lung (magenta) and liver (ochre), and cell lines (B). in B, osteosarcoma (U2OS, red), non-small cell lung cancer (H460 orange and A594 yellow), leukaemia (K562, pale green), breast cancer (MCF7 dark green and MDA-MB-231 turquoise green), colorectal carcinoma (DLD1 blue and HCT116 dark purple), cervical cancer (HeLa, pale purple), and HEK293T (brown).
